# Gliotransmission orchestrates neuronal type-specific axon regeneration

**DOI:** 10.1101/799635

**Authors:** Fei Wang, Kendra Takle Ruppell, Songlin Zhou, Yun Qu, Jiaxin Gong, Ye Shang, Yi Li, Yang Xiang

## Abstract

Why closely related neuronal types differ in their axon regenerative abilities remains elusive. Here, we demonstrate gliotransmission determines such a difference in *Drosophila* larval sensory neurons. Axotomy activates ensheathing glia, which signal to regenerative neurons through the gliotransmitter adenosine, to mount regenerative programs including neuronal activity and Ras. Surprisingly, ensheathing glia do not signal to non-regenerative neurons. Such neuronal type-specific responses to gliotransmission result from specific expression of adenosine receptors in regenerative neurons. Disrupting gliotransmission impedes regeneration of regenerative neurons. Strikingly, reconstitution of gliotransmission in non-regenerative neurons enables them to regenerate. Furthermore, activation of an adenosine receptor ortholog in adult mice promotes both regeneration and survival of retinal ganglion cells, uncovering a conserved pro-regenerative role of adenosine receptors. Our studies demonstrate gliotransmission as a novel mechanism by which glia instruct axon regeneration, with neuronal type-specificity, and suggest targeting purinergic signaling as a new strategy for mammalian central nervous system repair.

**HIGHLIGHTS:**

Ensheathing glia differentially interact with *Drosophila* sensory neuron types through gliotransmission
Gliotransmission mounts axon regenerative programs in selective neuronal types
Neuronal firing pattern but not overall excitability dictates axon regeneration outcome
Adenosine receptor activation in adult mice promotes both regeneration and survival of RGCs

## INTRODUCTION

Limited axon regeneration is a major hurdle to functional recovery after nervous system injury. Activation of the cell growth pathway has been shown to promote axon regeneration in different experimental systems, including that of adult central nervous system (CNS) neurons in mammals (He and Jin, 2016; Mahar and Cavalli, 2018). Surprisingly, the success of these manipulations differs drastically among closely related neuronal subtypes. For example, only alpha subtype retinal ganglion cells (RGCs) are capable of regenerating their axons after elevation of mammalian target of rapamycin (mTOR) activity, whereas other RGC subtypes fail to regenerate (Duan et al., 2015). Differences in regenerative ability among neuronal types is a general and conserved phenomenon that is observed in both the CNS and peripheral nervous system (PNS), as well as in other species including *Drosophila* and *C. elegans* (Canty et al., 2013; He and Jin, 2016; Song et al., 2012; Wu et al., 2007). However, how closely related neurons differ drastically in their regenerative abilities is poorly defined. Deciphering the underlying mechanisms is of great significance, as it will shed light on the fundamental question of how neurons respond to injury. Moreover, identifying the pro-regenerative programs from regenerative neurons and reactivating them in non-regenerative neurons could be a promising repair strategy.

Both neuronal intrinsic properties and their interactions with surrounding glial cells regulate axon regeneration (Yiu and He, 2006). The prevailing view is that axon regenerative competence is primarily determined by neuronal intrinsic properties, whereas glial cells modulate axon regeneration by forming glial scar and providing myelin-associated inhibitory factors (Yiu and He, 2006). However, a complete picture of how glial cells regulate axon regeneration has yet to emerge. Are there other types of glia-neuron crosstalk that could regulate axon regeneration? Could glia play any instructive role to establish axon regenerative programs? Gliotransmission is well characterized glia-neuron signaling by which glial cells rapidly communicate with neurons. Gliotransmission is initiated by glial Ca^2+^ signals that subsequently trigger release of small molecule transmitters known as gliotransmitters, including ATP and its metabolic product adenosine, glutamate, and D-serine. Gliotransmission has been primarily studied in the context of how astrocytes modulate synaptic strength (Bazargani and Attwell, 2016; Haydon, 2001; Lezmy et al., 2021; Ma et al., 2016; Pascual et al., 2005). However, whether and how gliotransmission could play a role in axon regeneration is yet to be determined.

Stimulating neuronal activity facilitates axon regeneration of mouse RGC axons (Li et al., 2016; Lim et al., 2016; Zhang et al., 2019). In these and other studies, the patterns of neuronal activity are not controlled, as such, it is unclear whether the firing pattern or general excitability increase underlies the pro-regenerative effect. Moreover, it is unknown whether regenerative and non-regenerative neurons could exhibit different neuronal activity. In addition, the Ca^2+^ transient immediately following axotomy is an early injury signal involved in diverse aspects of axon regeneration, including membrane resealing, growth cone formation, and activation of epigenetic programs (Cho et al., 2013; Ghosh-Roy et al., 2010; Mahar and Cavalli, 2018; Yan and Jin, 2012). However, it is unknown whether axotomy could elicit other Ca^2+^ signals to regulate regeneration.

Sensory neurons in the *Drosophila* larval PNS have emerged as a powerful experimental model for studying various aspects of neuronal regeneration (Song et al., 2019; Song et al., 2012; Thompson-Peer et al., 2016). These sensory neurons are divided into different subtypes based on their dendrite morphology and sensory functions (Grueber et al., 2003; Jan and Jan, 2001). Like mouse RGCs, different subtypes of *Drosophila* larval sensory neurons differ in their ability to regenerate their axons (Song et al., 2012). In this study, we exploit this genetically tractable system to dissect the mechanisms that control axon regeneration specificity, through parallel investigation of the regeneration-competent nociceptive neurons (aka class 4 dendritic arborization, or C4da) and the regeneration-incompetent touch-sensitive neurons (aka class 3 dendritic arborization, or C3da). We find that surrounding glial cells, which physically interact with each of larval sensory neurons by ensheathing their axon bundles (Yadav et al., 2019), actively respond to axotomy and play an instructive role in specifying the different axon regeneration abilities of C4da and C3da neurons through the gliotransmitter adenosine. We further show patterned neuronal activity, i.e., burst firing and Ca^2+^ spikes, is elicited in C4da, but not C3da, neurons to mediate the pro-regenerative action of gliotransmission. Finally, we reveal a conserved role of purinergic signaling in nervous system repair by showing that activation of the mammalian ortholog of the *Drosophila* adenosine receptor promotes both axon regeneration and survival of injured RGCs in adult mice.

## RESULTS

### Gliotransmission is essential for axon regeneration

Different types of sensory neurons in the *Drosophila* larval PNS reside in close proximity; their dendrites overlap, and their axons are bundled together and project to the ventral nerve cord (Figure S1A-S1C). These sensory neurons are surrounded by ensheathing glia that wrap the axon bundle (Figure S1B-S1C), forming a non-myelinating structure like the Remak bundle (Yadav et al., 2019; Yiu and He, 2006). Unless specified, we performed *in vivo* axotomy by focusing two-photon laser to a single axon, to minimize damages to neighboring axons and the ensheathing glia (Figure S1C). The distal axon degenerated after axotomy, and the proximal axon started to regrow at ∼24 hours post axotomy (hpa). To quantify regeneration, we measured axon length at 24 and 48 hpa to calculate a regeneration index that normalizes axon regrowth to larval body expansion during this period (Figure S1C, see also Star Methods). Consistent with a previous report (Song et al., 2012), we found nociceptive C4da neurons, which detect noxious heat, strong light, and irritant chemicals and are labeled by *ppk-CD4-tdGFP, ppk-CD4-tdTomato, ppk-EGFP* lines or the *ppk-Gal4/LexA* driver (Gu et al., 2019; Han et al., 2011; Hwang et al., 2007; Xiang et al., 2010), exhibited robust axon regeneration (Figure 1A-1D). In contrast, gentle touch-sensitive C3da neurons, labeled by *19-12-Gal4* in conjunction with *repo-Gal80*, or the *NompC*-*Gal4/LexA* driver (Xiang et al., 2010; Yan et al., 2013), failed to regenerate their axons (Figure 1A-1D).

**Figure 1.**
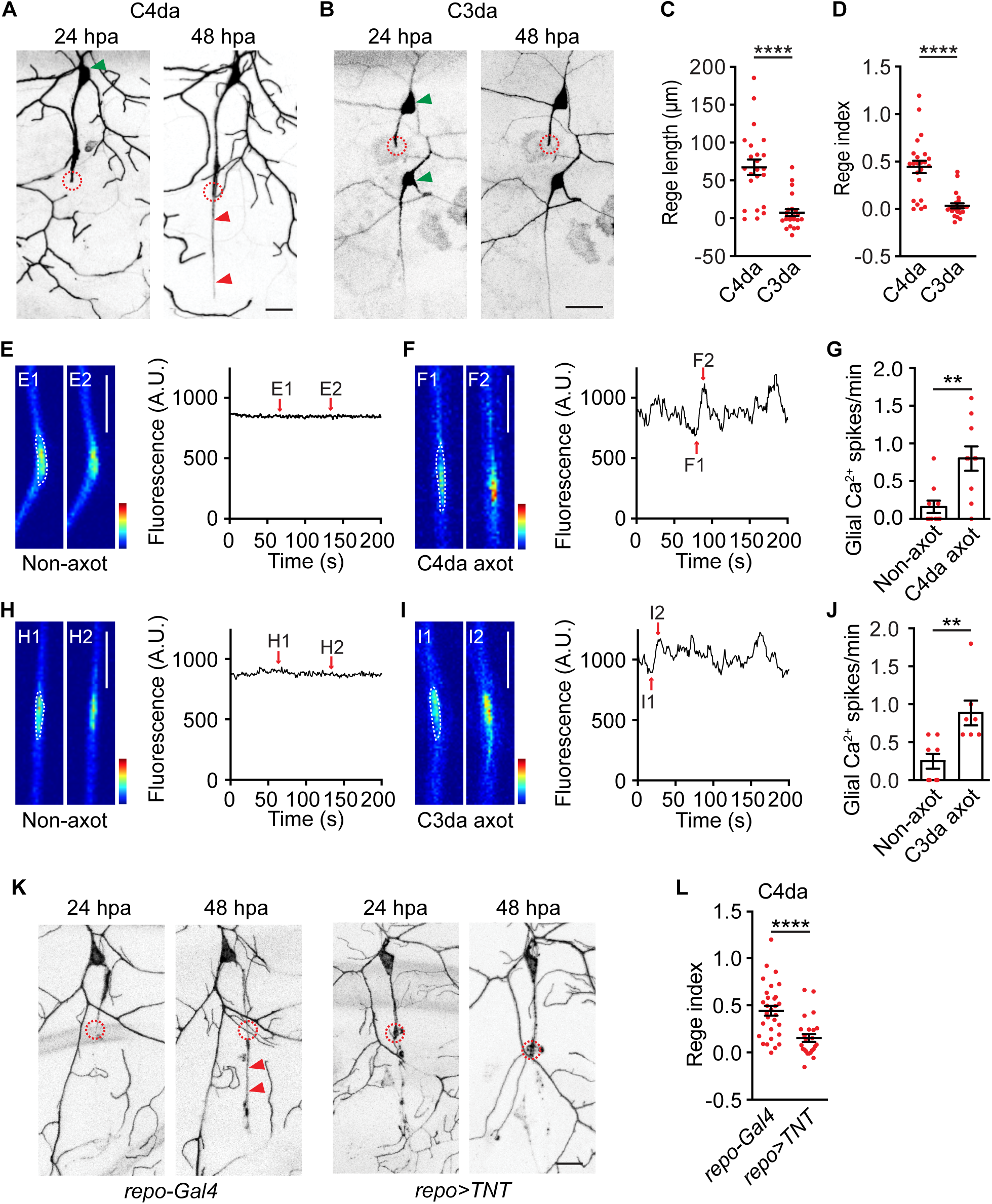
Gliotransmission is required for axon regeneration. (A-B) Representative images at 24 and 48 hours post axotomy (hpa) showing axon regeneration of a C4da neuron (A), but no axon regeneration of a C3da neuron (B). Circles indicated axotomy sites. Red arrowheads indicated the regrown axons, and green arrowheads indicated somata. Scale bar, 20 µm. Of the two C3da neurons, axotomy was performed on the dorsal neuron, ddaF (upper). (C-D) Quantification of axon regeneration length (C) and regeneration index (D). n=23 for C4da and n=22 for C3da neurons, respectively. Mann-Whitney test. (E-G) Axotomy of C4da neurons induced spontaneous glial Ca^2+^ spikes in ensheathing glia. (E) shows example images and traces of glial Ca^2+^ signals without axotomy (control). (F) shows example images and traces of glial Ca^2+^ signals at 24 hours after C4da axotomy. (G) shows quantification of glial Ca^2+^ spikes. Panels E1-E2 and F1-F2 showed individual image frames derived from the associated time-lapse GCaMP6s imaging traces. Glial Ca^2+^ spikes were quantified from regions indicated by white dashed lines. Scale bar, 20 µm. Color scale indicated dynamic range from 0 to 2,500. n=10 and 10. Mann-Whitney test. (H-J) Axotomy of C3da neurons induced spontaneous glial Ca^2+^ spikes in ensheathing glia. (H) shows example images and traces of glial Ca^2+^ signals without axotomy (control). (I) shows example images and traces of glial Ca^2+^ signals at 24 hours after C3da axotomy. (J) shows quantification of glial Ca^2+^ spikes. Panels H1-H2 and I1-I2 showed individual image frames derived from the associated time-lapse GCaMP6s imaging traces. Glial Ca^2+^ spikes were quantified from regions indicated by white dashed lines. Scale bar, 20 µm. Color scale indicated dynamic range from 0 to 3,000. n=8 and 7. Mann-Whitney test. (K-L) Representative images (K) and regeneration index (L) of axon regeneration of C4da neurons from control larvae (*repo-Gal4*) and larvae expressing TNT in glia (*repo-Gal4>UAS-TNT*). Circles indicated axotomy sites and red arrowheads showed regenerated axons. n=30 and 24. Mann-Whitney test. Scale bar, 20 µm. * *P*<0.05, ** *P*<0.01, *** *P*<0.001, **** *P*<0.0001. n. s., not significant. Data are represented as mean ± S.E.M.. See also Figure S1 and S2.

To assess the role of ensheathing glia in axon regeneration, we increased the laser scanning area to damage both axon and its associated ensheathing glia, a paradigm we termed as glial bundle cut. Although morphological integrity of ensheathing glia was intact after standard axotomy, it was disrupted after glial bundle cut (Figure S2A). Strikingly, damaging glia drastically reduced axon regeneration of C4da neurons (Figure S2A-S2B). These results suggest a possibility that glia-derived signals could regulate axon regeneration. Next, we performed GCaMP imaging to determine how ensheathing glia respond to axotomy. We found that Ca^2+^ levels in ensheathing glia were stable without axotomy, however, axotomy of C4da neurons elicited spontaneous Ca^2+^ oscillations in ensheathing glia that we termed as glial Ca^2+^ spikes (Figure 1E-1G). Similar glial Ca^2+^ spikes were observed after axotomy of C3da neurons (Figure 1H-1J), indicating that ensheathing glia are activated by axotomy regardless of neuronal types being injured. A major effector of glial Ca^2+^ elevation is release of gliotransmitters, a process known as gliotransmission (Haydon, 2001). Despite extensive studies on the roles of glial scar and myelin-associated factors in axon regeneration (Yiu and He, 2006), whether gliotransmission could play a role in axon regeneration is surprisingly unknown. Gliotransmission is disrupted after functional blockage of soluble N-ethylmaleimide-sensitive factor attachment protein receptor (SNARE) (Agulhon et al., 2008; Bezzi et al., 2004; Jourdain et al., 2007; Panatier et al., 2011; Pascual et al., 2005; Perea and Araque, 2007). We thereby perturbed the SNARE function in glia to determine the role of gliotransmission on axon regeneration. We found that glial expression of tetanus toxin (TNT), a neurotoxin that cleaves the vesicular SNARE synaptobrevin (Sweeney et al., 1995), significantly impaired axon regeneration of C4da neurons (Figure 1K-1L). Together, our results indicate that ensheathing glia are activated by axotomy and critically control C4da neuron axon regeneration through gliotransmission.

### Gliotransmission evokes neuronal type-specific responses through gliotransmitter adenosine

How might gliotransmission regulate axon regeneration? To address this question, we first activated ensheathing glia to determine the consequence on sensory neurons. TrpA1-B (i.e., TrpA1 alternative isoform B) is a Ca^2+^-permeable cationic channel with high sensitivity to warmth (Gu et al., 2019). Here, we expressed TrpA1-B in glial cells and heated dissected larvae to ∼33°C, temperature that is sufficient to activate TrpA1-B but cannot activate C4da neurons (Gu et al., 2019; Xiang et al., 2010). As expected, ensheathing glia expressing TrpA1-B, but not control ensheathing glia, responded to heat with increased Ca^2+^ levels (Figure S2D-S2E). Gliotransmitters have been shown to act on their neuronal receptors to regulate neuronal excitability (Haydon, 2001). We therefore determined whether ensheathing glial could modulate sensory neuron activity. GCaMP imaging of sensory neurons showed that ensheathing glial stimulation elicited robust Ca^2+^ responses in C4da neurons, but surprisingly had little effect in C3da neurons (Figure 2A-2B, Video S1 and S2). This C4da neuron response was abolished after glial expression of TNT (Figure 2B), indicating that ensheathing glia activate C4da neurons through gliotransmission. To corroborate this finding, we stimulated ensheathing glia with optogenetics, which offers better temporal control over thermogenetics. Consistently, we found light stimulation of CsChrimson-expressing glia activated C4da neurons (Figure S2F-S2G). The short latency between light stimulation and neuronal responses further indicates that gliotransmission can quickly modulate neuronal activity. Together, our results demonstrate that ensheathing glia specifically activate regenerative C4da, but not non-regenerative C3da, neurons, through gliotransmission.

**Figure 2.**
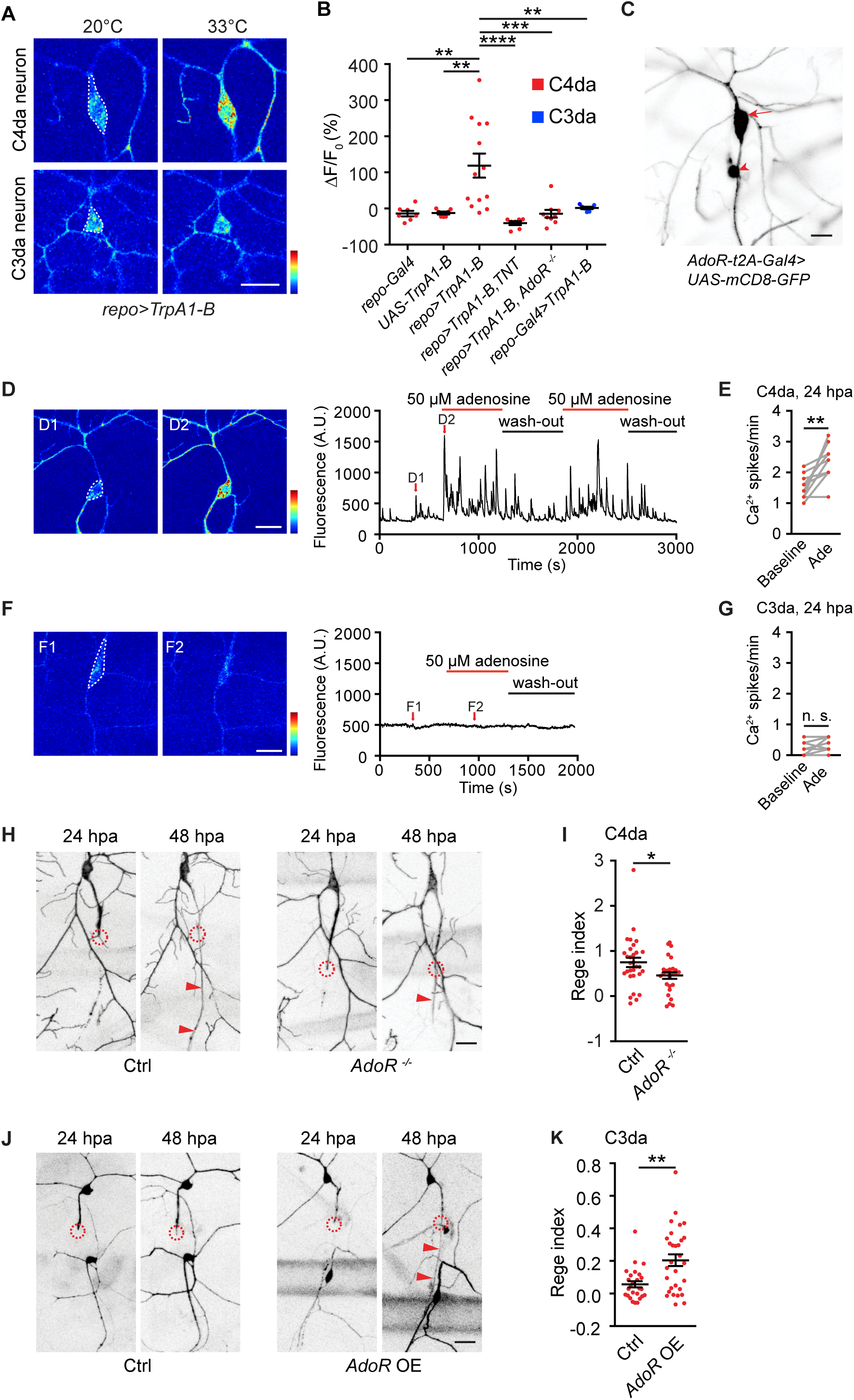
Gliotransmission determines axon regeneration differences of neuronal types. (A) Representative Ca^2+^ response of C4da and C3da neurons in response to acute thermogenetic stimulation of TrpA1-B-expressing glia. TrpA1-B is activated at 33°C but not 20°C. Color scale indicated dynamic range from 0 to 2,500. Scale bar, 20 µm. (B) Quantifications of (A) showing peak GCaMP6s signals of C4da and C3da neurons in response to glial thermogenetic stimulation. C4da, but not C3da neurons were activated by glial stimulation. This C4da neuron response was abolished in *AdoR* mutant larvae, and in larvae in which TNT was co-expressed in glia. Signals were quantified from somata as indicated by white dashed lines in (A). n=7, 7, 13, 7, 9, 7. One-way ANOVA followed by Tukey’s test. (C) Representative imaging of a 3^rd^ instar larva bearing *AdoR-t2A-Gal4*>*UAS-mCD8-GFP* showed AdoR expression in the C4da neuron (arrow), but not in C3da neurons or glia. In addition, AdoR is found in another sensory neuron likely to be dmd (arrowhead). Scale bar, 20 µm. (D-E) Representative images (D) and summary results (E) of Ca^2+^ responses of axotomized C4da neurons to adenosine application. D1-D2 showed individual image frames derived from the associated time-lapse GCaMP6s imaging traces shown in the right panel. Scale bar, 20 µm. Color scale indicated dynamic range from 0 to 3,000. In this example, adenosine was perfused twice followed by washout. The Ca^2+^ spike frequency of axotomized C4da neurons was quantified from the somata (indicated by white dashed lines in (D)) before and during adenosine application. n=10. Two-tailed paired *t*-test. (F-G) Representative images (F) and summary results (G) of Ca^2+^ responses of axotomized C3da neurons to adenosine application. F1-F2 showed individual image frames derived from the associated time-lapse GCaMP6s imaging traces shown in the right panel. Scale bar, 20 µm. Color scale indicated dynamic range from 0 to 3,000. The Ca^2+^ spike frequency of axotomized C3da neurons was quantified from the somata (indicated by white dashed lines in (F)) before and during adenosine application. n=8. Two-tailed paired *t*-test. (H-I) Representative images of axon regeneration (H) and regeneration index (I) of C4da neurons from wildtype larvae (control) and larvae bearing *AdoR* deletion mutant. Circles indicated axotomy sites and red arrowheads showed regenerated axons. n=30 and 27. Mann-Whitney test. Scale bar, 20 µm. (J-K) Representative images of axon regeneration (J) and regeneration index (K) of C3da neurons from control larvae (*19-12-Gal4*) and larvae bearing AdoR overexpression (OE) in C3da neurons (*19-12-Gal4>UAS-AdoR*). Circles indicated axotomy sites and red arrowheads showed regenerated axons. n=26 and 30. Mann-Whitney test. Scale bar, 20 µm. * *P*<0.05, ** *P*<0.01, *** *P*<0.001, **** *P*<0.0001. n. s., not significant. Data are represented as mean ± S.E.M.. See also Figure S2, S3, Video S1 and S2.

In both mammals and *Drosophila*, ATP and its metabolic product adenosine are gliotransmitters (Lezmy et al., 2021; Ma et al., 2016; Pascual et al., 2005). The *Drosophila* genome encodes a single purinergic receptor, adenosine receptor (AdoR), a G protein-coupled receptor (GPCR) family member that couples to Gαs and elicits Ca^2+^ elevation in response to adenosine stimulation (Dolezelova et al., 2007; Ma et al., 2016; Xu et al., 2020). We found C4da neuron responses elicited by ensheathing glial stimulation was abolished in AdoR mutant larvae (Figure 2B), indicating that adenosine is the gliotransmitter that mediates glia-to-neuron signaling.

Next, we acutely applied adenosine to determine the effect on axotomized sensory neurons. We found adenosine increased the frequency of spontaneous Ca^2+^ spikes of axotomized C4da neurons (Figure 2D and 2E, Ca^2+^ spikes in C4da neurons will be discussed later), in contrast, axotomized C3da neurons did not respond to adenosine (Figure 2F-2G). Moreover, glutamate and D-serine, two other gliotransmitters (Haydon, 2001), had no effects on axotomized C4da neurons (Figure S3A-S3D). These results indicate that gliotransmission differentially modulates C4da and C3da neuron activity through adenosine.

### Reconstitution of gliotransmission responses in C3da neurons enables them to regenerate

To assess how C4da and C3da neurons differentially responded to gliotransmission and adenosine, we examined AdoR expression using the recently generated AdoR-t2A-Gal4 reporter fly strain (Deng et al., 2019). In this knock-in strain, t2A-Gal4 is inserted into the AdoR locus immediately before the stop codon so that AdoR and Gal4 are produced as a single peptide during translation but are later separated *via* t2A-mediated self-cleavage (Deng et al., 2019).

Hence, Gal4 activity in AdoR-t2A-Gal4 flies reports the endogenous AdoR expression. A similar strategy has been successfully used to report *in vivo* expression of other ion channels and receptors in *Drosophila* (Deng et al., 2019; Gu et al., 2019). By crossing AdoR-t2A-Gal4 to a reporter line, we found intriguingly that AdoR was expressed in C4da neurons, but not in C3da neurons or ensheathing glia (Figure 2C). This neuron type-specific AdoR expression provides a molecular explanation of neuronal type-specific responses to gliotransmission and adenosine.

We then assessed the role of AdoR in axon regeneration (Deng et al., 2019). We found that axon regeneration of C4da neurons was impaired in AdoR mutant larvae (Figure 2H-2I). Therefore, disrupting gliotransmission, either by glial expression of TNT or by AdoR mutation causes impaired axon regeneration. Glial Ca^2+^ spikes were elicited after axotomy of either C4da or C3da neurons (Figure 1E-1J), suggesting that enshething glia could release ATP/adenosine after axotomy of either neuronal type. We therefore hypothesized that differential regenerative abilities of C4da and C3da neurons are determined by their distinct responses to gliotransmission. To test this hypothesis, we reconstituted gliotransmission responses by ectopic expression of AdoR in C3da neurons. As positive control, we showed that adenosine application activated AdoR-expressing C3da neurons, but not control C3da neurons (Figure S3E-S3F). Strikingly, while control C3da neurons failed to regenerate after laser axotomy, AdoR-expressing C3da neurons displayed robust axon regeneration (Figure 2J-2K). Because gliotransmission is necessary for C4da neuron axon regeneration and is sufficient to induce C3da neuron axon regeneration, our results reveal an instructive role of gliotransmission in promoting axon regeneration. These results further demonstrate gliotransmission as a novel mechanism by which glia control axon regeneration with neuronal type-specificity.

### Gliotransmission elicits Ca^2+^ spikes and burst firing as regenerative programs in C4da, but not C3da, neurons

What are the axon regenerative programs in neurons that are activated by gliotransmission? We reasoned these regenerative programs should meet the following three criteria; i) they are active in axotomized C4da neurons and depend on gliotransmission, ii) they are inactive in axotomized C3da neurons but are activated by AdoR expression, and iii) disrupting these programs would impair C4da neuron axon regeneration and reactivating these programs would enable C3da neurons to regenerate. We hypothesize that neuronal Ca^2+^ signals could be one such regenerative program, based on findings that both glial activation and adenosine application elevated Ca^2+^ signals in C4da, but not C3da, neurons (Figure 2A-2B and 2D-2G). We tested this hypothesis by performing GCaMP6 imaging (Chen et al., 2013; Gu et al., 2019), to determine intrinsic Ca^2+^ responses of axotomized and non-axotomized sensory neurons in the same larvae at 24 hpa. We found that axotomized C4da neurons exhibited spontaneous Ca^2+^ spikes that were synchronized across soma, axon and dendrites, in contrast, Ca^2+^ spikes were seldom detected in non-axotomized C4da neurons (Figure 3A-3B, 3E, S4A-S4E, Video S3 and S4). This Ca^2+^ spike of axotomized C4da neurons were abolished after removal of extracellular Ca^2+^ (Figure S4F), indicating the requirement of Ca^2+^ influx. Next, we determined the time course of Ca^2+^ spikes by imaging C4da neurons at different time points after axotomy. We found that post axotomy, Ca^2+^ spikes had delayed onset and gradually increased over time; their frequency was initially low in the first few hours, reached a plateau at ∼18 hpa and sustained afterwards (Figure 3G). Therefore, Ca^2+^ spikes preceded and persisted throughout the time window of axon regeneration of C4da neurons, which started at ∼24 hpa (Figure S1C) (Song et al., 2012). This time window raises the possibility that Ca^2+^ spikes could regulate axon regeneration. In contrast to C4da neurons, we found Ca^2+^ spikes in neither non-axotomized nor axotomized C3da neurons (Figure 3C-3D and 3F). Moreover, Ca^2+^ spikes do not reflect a general injury response, as Ca^2+^ spikes were not elicited after severing dendrites of C4da neurons or after induction of systemic inflammatory responses by exposing larvae to UV-C light (Figure 3E) (Babcock et al., 2009). Together, these results indicate that spontaneous Ca^2+^ spikes are neuronal type-specific and axotomy specific.

**Figure 3.**
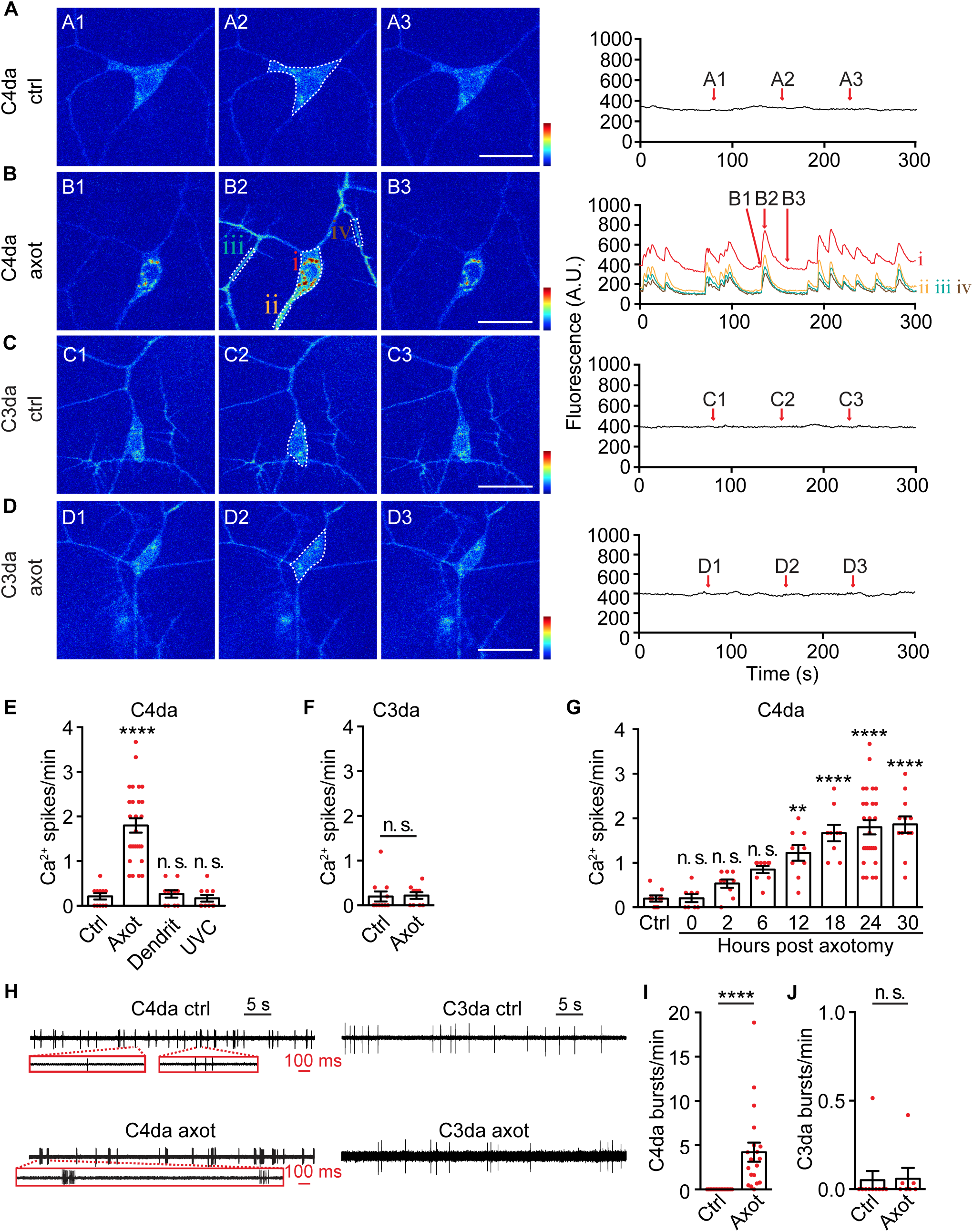
Axotomy elicits Ca^2+^ spikes and burst firing in C4da, but not C3da, neurons. (A-D) Representative spontaneous Ca^2+^ signals, measured by GCaMP6s imaging, from control (ctrl, non-axotomized) and axotomized (axot) C4da neurons (A-B), and from control (ctrl, non-axotomized) and axotomized (axot) C3da neurons (C-D). Left three panels (e.g., A1-A3) showed individual image frames at given time points indicated in the time-lapse GCaMP6s imaging traces (right panel). For traces shown in (A), (C), and (D), GCaMP6s signals from cell body as indicated by dashed lines were shown. For traces shown in (B), GCaMP6s signals from cell body (i), axon (ii) and dendrites (iii and iv) as indicated by white dashed lines were shown. Color scale indicated dynamic range from 0 to 1,800. Scale bar, 20 µm. (E) Average frequency of spontaneous Ca^2+^ spikes (spikes/min) of C4da neurons in response to axotomy (axot, 24 hpa), dendritomy (dendrit, 24 hours after lesion) and 256nm UV-C light exposure (UVC, 24 hours after exposure). n=11, 27, 10 and 10. One-way ANOVA followed by Bonferroni’s tests between control and treated groups. (F) Average frequency of spontaneous Ca^2+^ spikes of control (ctrl, non-axotomized) and axotomized (axot) C3da neurons at 24 hpa. n=11 and 9. Mann-Whitney test. (G) Average frequency of spontaneous Ca^2+^ spikes of C4da neurons at different time points after axotomy. n=9, 8, 9, 9, 9, 9, 27 and 12 for each time points. One-way ANOVA followed by Dunnett’s tests between control (ctrl, non-axotomized) and other axotomized groups. (H) Representative traces of extracellular recording showing spontaneous firing of control (ctrl, i.e., non-axotomized) and axotomized (axot) C4da and C3da neurons. Zoom-in views (red boxes) show representative burst firing events in axotomized C4da neurons. (I) The average frequency of burst firing events (bursts/min) of C4da neurons. n=17 for control C4da neurons and n=19 for axotomized C4da neurons. Mann-Whitney test. (J) The average frequency of burst firing events (bursts/min) of C3da neurons. n=10 for control C3da neurons and n=7 for axotomized C3da neurons. Mann-Whitney test. * *P*<0.05, ** *P*<0.01, *** *P*<0.001, **** *P*<0.0001. n. s., not significant. Data are represented as mean ± S.E.M.. See also Figure S4, S5, Video S3, S4 and S5.

Ca^2+^ transients immediately following axotomy regulate early events of axon regeneration such as membrane resealing and growth cone formation (Cho et al., 2013; Ghosh-Roy et al., 2010; Mahar and Cavalli, 2018). Unlike Ca^2+^ spikes that are specific to C4da neurons, immediate Ca^2+^ transients were elicited in both C3da and C4da neurons after axotomy (Figure S4G-S4H), suggesting immediate Ca^2+^ transients are unlikely to underlie differential regenerative abilities of C4da and C3da neurons. These results also suggest immediate Ca^2+^ transients and Ca^2+^ spikes are likely generated by distinct mechanisms. Supporting this notion, Ca^2+^ spikes emerged hours after axotomy and persisted throughout the axon regeneration time window, whereas immediate Ca^2+^ transients appeared immediately after axotomy and were transient. Therefore, we propose spontaneous Ca^2+^ spikes as a new injury signal.

To determine whether spontaneous Ca^2+^ spikes reflect axotomy-induced neuronal activity changes, we performed extracellular recordings on axotomized and non-axotomized neurons in the same larvae at 24 hpa. We found axotomy transformed the firing pattern of C4da neurons by eliciting spontaneous burst firing events, where rapid action potential sequences were discharged intermittently (Figure 3H, see Star Methods for definition of burst firing events). These bursting firing events were frequently observed in axotomized C4da neurons (18 out of 19) but barely in non-axotomized C4da neurons (1 out of 17) (Figure S5C and S5E). Moreover, the frequency of burst firing events, and the percentage of total action potentials found in the burst firing events, were significantly increased in axotomized C4da neurons when compared to non-axotomized C4da neurons (Figure 3I and Figure S5F). However, the overall firing frequency was similar in axotomized and non-axotomized C4da neurons (Figure S5A). Therefore, axotomy did not significantly alter C4da neuron excitability, but rather clustered action potentials into burst firing events. In contrast to C4da neurons, axotomy did not elicit burst firing or otherwise alter the excitability of C3da neurons (Figure 3H, 3J, Figure S5B, S5D-S5E and S5G). To further assess the relationship between burst firing and Ca^2+^ spikes, we performed simultaneous recording and Ca^2+^ imaging on axotomized C4da neurons. We found a high temporal correlation between burst firing and Ca^2+^ spikes (Figure S4I and Video S5), suggesting that Ca^2+^ spikes result from burst firing-induced membrane depolarization. Hence, Ca^2+^ spikes can serve as a surrogate for reporting burst firing events.

After showing that axotomy induced spontaneous Ca^2+^ spikes specifically in C4da neurons, we set out to determine whether Ca^2+^ spikes depend on gliotransmission. We found Ca^2+^ spikes in axotomized C4da neurons were significantly reduced after blocking gliotransmission by TNT (Figure 4A-4C), and in AdoR mutants (Figure 4D-4F). Consistently, damaging glia by ‘glial bundle cut’ abolished burst firing in axotomized C4da neurons (Fig. S2C). Next, we determined whether reconstitution of gliotransmission responses in C3da neurons, which was sufficient to elicit axon regeneration (Fig. 2J-2K), could elicit spontaneous Ca^2+^ spikes in axotomized C3da neurons. Strikingly, although no Ca^2+^ spikes were detected in control axotomized C3da neurons, AdoR-expressing C3da neurons exhibited robust Ca^2+^ spikes after axotomy (Figure 4G-4I). These Ca^2+^ spikes were synchronized across soma and dendrites of C3da neurons, similar to those found in axotomized C4da neurons (Figure 3B). Moreover, induced Ca^2+^ spikes in C3da neurons required axotomy, as non-axotomized AdoR-expressing C3da neurons did not show Ca^2+^ spikes (Figure S3G-S3H). Together, our results indicate that Ca^2+^ spikes in axotomized C4da neurons depend on gliotransmission, and that reconstituted gliotransmission is sufficient to induce Ca^2+^ spikes in axotomized C3da neurons.

**Figure 4.**
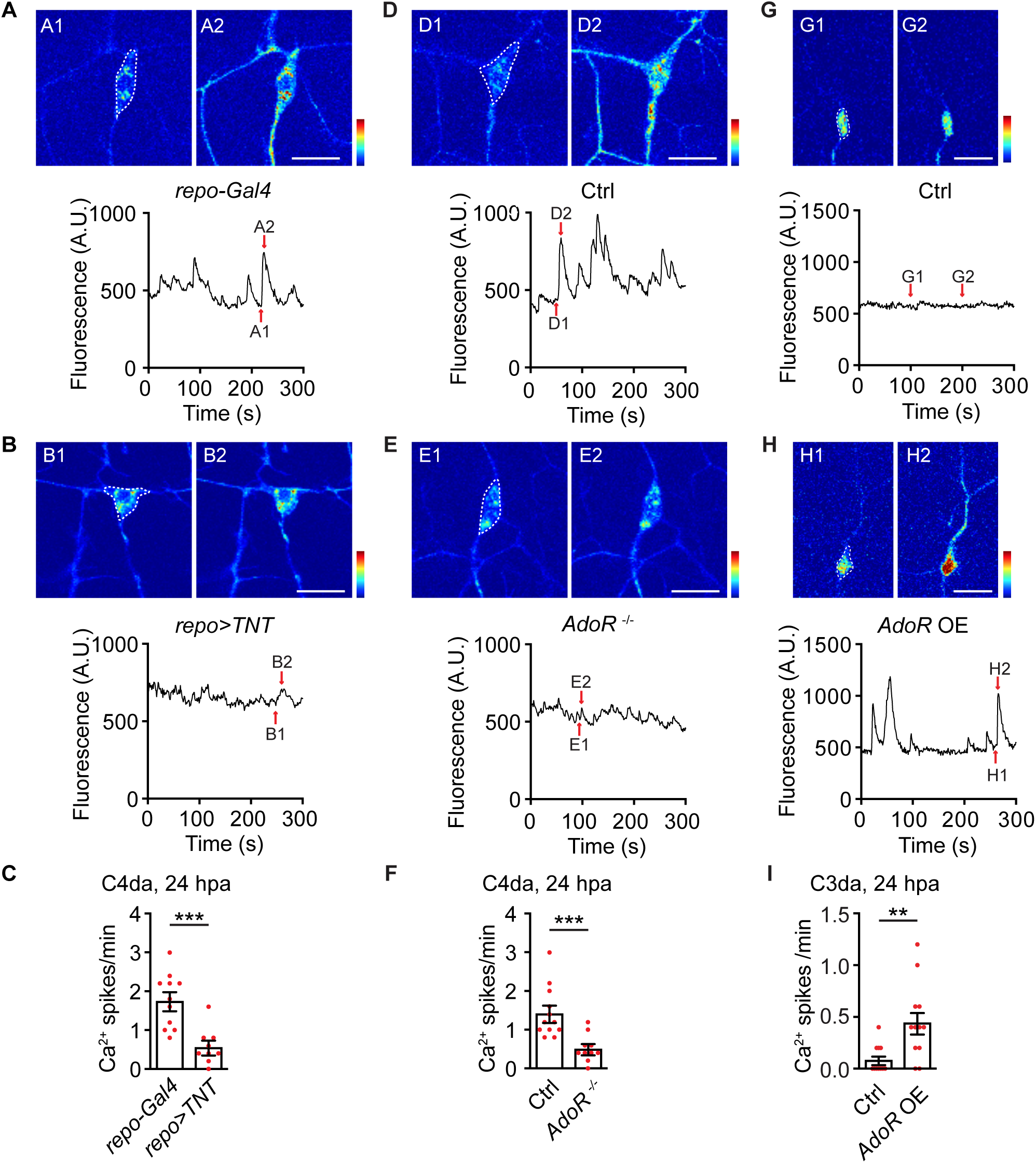
Gliotransmission is necessary for C4da neuron Ca^2+^ spikes and reconstitution of gliotransmission is sufficient to induce Ca^2+^ spikes in C3da neurons. (A-B) Representative spontaneous Ca^2+^ signals, measured by GCaMP6s imaging, from axotomized C4da neurons (*repo-Gal4*, panel A) and axotomized C4da neurons with glia overexpressing TNT (*repo-Gal4>UAS-TNT*, panel B) at 24 hpa. Top panels (e.g., A1-A2) showed individual image frames at given time points indicated in the time-lapse GCaMP6s imaging traces from white dashed lines (bottom panels). Color scale indicated dynamic range from 0 to 2,500. Scale bar, 20 µm. (C) Glial expression of TNT reduced the frequency of spontaneous Ca^2+^ spikes in axotomized C4da neurons. n=11 and 9. Two-tailed unpaired *t*-test. (D-E) Representative spontaneous Ca^2+^ signals, measured by GCaMP6s imaging, from axotomized C4da neurons of wildtype (control) larvae (D) and larvae bearing *AdoR* deletion mutant (E) at 24 hpa. Top panels (e.g., D1-D2) showed individual image frames at given time points indicated in the time-lapse GCaMP6s imaging traces from white dashed lines (bottom panels). Color scale indicated dynamic range from 0 to 2,500. Scale bar, 20 µm. (F) Quantification of the frequency of spontaneous Ca^2+^ spikes of axotomized C4da neurons from wildtype (control) and *AdoR* deletion mutant larvae. n=12 and 10. Two-tailed unpaired *t*-test. (G-I) Representative examples (G, H) and summary (D) of spontaneous Ca^2+^ spikes of axotomized C3da neurons from control larvae (*19-12-Gal4*>*UAS-GCaMP6s*) (G) and larvae bearing AdoR overexpression (OE) in C3da neurons (*19-12-Gal4*>*UAS-AdoR, GCaMP6s*) (H). G1-G2 and H1-H2 show individual image frames derived from the associated time-lapse GCaMP6s imaging traces shown in the bottom panels. Color scale indicated dynamic range from 50 to 1,400. Scale bar, 20 µm. The Ca^2+^ spike frequency was quantified from the somata of C3da neurons as indicated by white dashed lines. n=11 and 12. Two-tailed unpaired *t*-test. * *P*<0.05, ** *P*<0.01, *** *P*<0.001, **** *P*<0.0001. n. s., not significant. Data are represented as mean ± S.E.M.. See also Figure S3.

### Firing pattern but not overall excitability dictates axon regeneration outcome

To assess whether neuronal Ca^2+^ spikes are required for axon regeneration, we expressed the inward rectifying potassium channel Kir2.1 in C4da neurons (Yang et al., 2014). We found Kir2.1 reduced frequency of spontaneous Ca^2+^ spikes (Figure 5A), as well as axon regeneration of axotomized C4da neurons (Figure 5B-5C). To further investigate the role of Ca^2+^ signals, we expressed parvalbumin, a vertebrate specific Ca^2+^ binding protein that can sequester Ca^2+^ in *Drosophila* tissues (Weavers et al., 2016). Like Kir2.1, parvalbumin expression in C4da neurons significantly reduced both Ca^2+^ spikes and axon regeneration (Figure 5A-5C). These results indicate that spontaneous neuronal activity is necessary for axon regeneration of C4da neurons.

**Figure 5.**
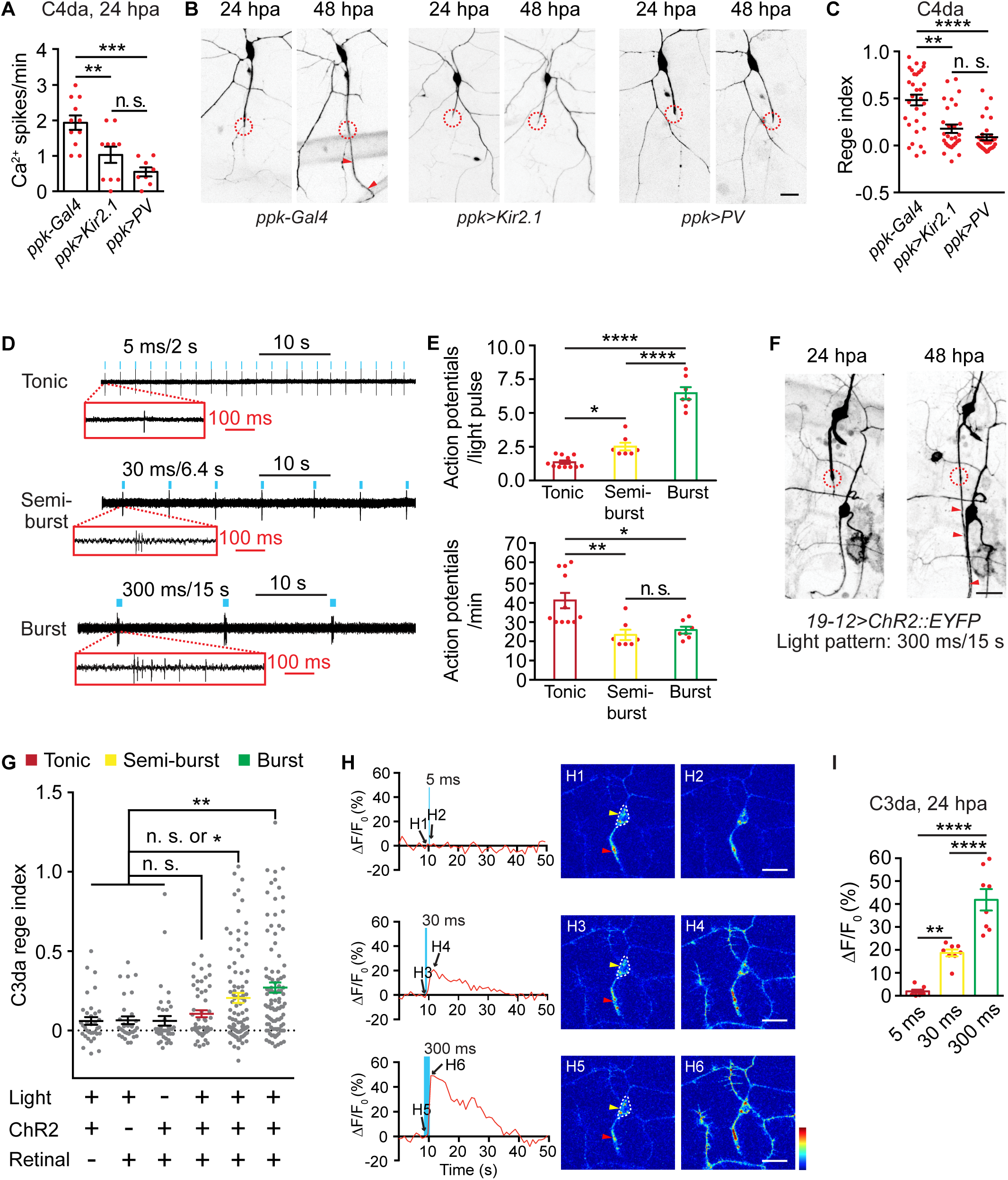
Firing pattern instead of excitability determines axon regeneration outcome. (A) Quantification of the frequency of spontaneous Ca^2+^ spikes of axotomized C4da neurons bearing *ppk-Gal4* (control), or axotomized C4da neurons that express Kir2.1 driven by *ppk-Gal4* (*ppk-Gal4>UAS-Kir2.1*) or axotomized C4da neurons that express parvalbumin (PV) driven by *ppk-Gal4* (*ppk-Gal4>UAS-PV*). n=11, 10 and 8. One-way ANOVA followed by Bonferroni’s tests. (B) Representative images of axon regeneration of control, Kir2.1-expressing and PV-expressing C4da neurons. Circles indicated axotomy sites and red arrowheads showed regenerated axons. Scale bar, 20 µm. (C) Quantification of regeneration index of control, Kir2.1-expressing and PV-expressing C4da neurons. n=31, 28 and 29. Kruskal-Wallis test followed by Dunn’s tests. (D) Representative extracellular recordings of C3da neurons showing three different firing patterns evoked by optogenetics. Delivery of 470 nm, 1.4 mW/mm^2^ blue light was marked by blue dots (not to scale). Scale bar, 10 s. Zoom-in traces show spike patterns. Scale bar, 100 ms. (E) Upper panel, quantification of action potential numbers in response to a single optogenetic light pulse stimulation. n=11, 7 and 7; One-way ANOVA followed by Bonferroni’s tests. Lower panel, quantification of average action potential numbers per minute in response to different optogentic stimulation patterns. n=11, 7 and 7; One-way ANOVA followed by Bonferroni’s tests. (F) Representative images showing axon regeneration of C3da neurons elicited by optogenetics-induced burst firing. Circles indicated axotomy sites and red arrowheads showed the regenerated axon. Scale bar, 20 µm. (G) Quantification of axon regeneration of C3da neurons under different optogenetic stimulation conditions: three control groups (no retinal, n=35; no ChR2, n=32; no light n=39) and three optogenetic stimulation groups (tonic firing, n=51; semi-burst firing, n=99; burst firing, n=97). Kruskal-Wallis test followed by Dunn’s tests. (H) Representative Ca^2+^ responses of axotomized C3da neurons at 24 hpa to a single light pulse that is used to induce tonic, semi-burst and burst firing. Left panels showed corresponding time-lapse GCaMP6s imaging traces from soma, indicated by white dashed lines in right panels. Blue bars marked the delivery of 5 ms, 30 ms and 300 ms 470 nm, 1.4 mW/mm^2^ blue light for tonic, semi-burst and burst firing, respectively. Right panels (e.g., H1-H2) showed GCaMP6s signals before and immediately after a single light pulse stimulation. Red and yellow arrowheads indicated axons and somata, respectively. Scale bar, 20 µm. Color scale indicated dynamic range from 0 to 1,500. (I) Summary results of (H) by quantifying peak GCaMP6s responses from somata of axotomized C3da neurons. n=8, 8 and 8. One-way ANOVA followed by Bonferroni’s tests. * *P*<0.05, ** *P*<0.01, *** *P*<0.001, **** *P*<0.0001. n. s., not significant. Data are represented as mean ± S.E.M.. See also Figure S6.

Previous studies indicate that stimulating neuronal activity promotes axon regeneration (Li et al., 2016; Lim et al., 2016; Zhang et al., 2019). However, it is yet unknown whether neuronal excitability or a specific firing pattern is most responsible for the observed effect. Because axotomy induced burst firing in C4da neurons (Figure 5A-5C), we hypothesized that a specific firing pattern is critical. To test this hypothesis, we exploited optogenetics to stimulate axotomized C3da neurons in behaving larvae with three distinct firing patterns including burst firing, semi-burst firing, and tonic firing, while maintaining the total action potential numbers comparable (Figure 5D-5E). Burst firing stimulation was achieved by ‘replaying’ spontaneous activity of axotomized C4da neurons onto axotomized C3da neurons. To this goal, we first determined the burst firing characteristics of axotomized C4da neurons (7.3 action potential/burst, 4.2 bursts/minute and 40.8 ms inter-spike interval, Figure 3I and Figure S6A-S6B). Next, we expressed light-sensitive channelrhodopsin2 (ChR2) in C3da neurons and recorded their responses to 470 nm blue light (Fenno et al., 2011). By varying intensity, duration, and frequency of light pulses, we found a specific stimulation pattern that elicited burst firing events resembling those of axotomized C4da neurons (1.4 mW/mm^2^, 300 ms duration for every 15 s, see ‘burst’ in Figure 5D-5E, Figure S6A-S6B). Because Ca^2+^ spikes appeared at ∼6 hpa in C4da neurons (Figure 3G), we stimulated C3da neurons starting at ∼6 hpa and lasting throughout the experiment (Figure S6C-S6D). Intriguingly, burst firing stimulation elicited robust axon regeneration of C3da neurons (Figure 5F and ‘burst’ in Figure 5G), when compared to control groups where ChR2, light, or retinal was not supplied (Figure 5G). The magnitude of C3da neuron axon regeneration induced by burst firing was comparable to that after AdoR expression (regeneration index: 0.205±0.0361, n=30, AdoR expression *versus* 0.271±0.0319, n=97, optogenetics, Mann-Whitney test, *P*=0.779). We next stimulated C3da neurons with tonic firing and semi-burst firing, which elicited less action potential numbers per light pulse than burst firing, but nevertheless higher (tonic) or comparable (semi-burst) numbers of total action potentials per minute (Figure 5D-5E). We found that tonic firing was the least effective, whereas semi-burst firing was intermediate in promoting C3da neuron axon regeneration (Figure 5G). These results provided direct evidence the neuronal activity pattern, rather than overall excitability determines axon regeneration strength. We further assessed whether different activity patterns could evoke differential Ca^2+^ responses. Indeed, burst firing evoked the strongest Ca^2+^ spikes in axotomized C3da neurons (Figure 5H-5I). These Ca^2+^ spikes were synchronized across the cell body, axon and dendrites (Figure 5H), similar to those observed in axotomized C4da neurons (Figure 3B). In contrast, semi-burst firing evoked weak Ca^2+^ spikes, while tonic firing produced no overt Ca^2+^ responses (Figure 5H-5I). Therefore, burst firing evoked the strongest axon regeneration, likely through Ca^2+^ spike induction. Altogether, our data indicate spontaneous Ca^2+^ spikes and burst firing as the neuronal type-specific axon regenerative program downstream of gliotransmission by satisfying the three aforementioned criteria. Moreover, we provide the first evidence that firing pattern instead of overall excitability dictates axon regeneration outcome.

### Neuronal type-specific Ras activity determines axon regeneration specificity

To investigate how Ca^2+^ spikes determine axon regeneration, we focused on Ras GTPase as a potential downstream effector based on the findings that intracellular Ca^2+^ activates Ras and that mitogen-activated protein kinase (MAPK) pathways regulate axon regeneration in both mammals and flies (Hollis et al., 2009; O’Donovan et al., 2014; Perlson et al., 2005; Rosen et al., 1994; Wang et al., 2020). We found that reducing Ras activity in C4da neurons, by expressing the *Drosophila* or mouse version of dominant negative Ras (Ras^DN^), significantly impaired axon regeneration (Figure 6A-6B). Ras activity is reported by diphospho-extracellular signal–regulated kinase (dpERK) (Gabay et al., 1997; Xu et al., 2020). We performed immunostaining and found that dpERK levels were low in both non-axotomized C4da and C3da neurons, however, axotomy significantly increased dpERK in C4da neurons, but not in C3da neurons (Figure 6E-6H). These findings indicate that axotomy causes neuronal type-specific Ras activation. To test whether Ras activation is sufficient for axon regeneration, we expressed constitutively active Ras (Ras^CA^) in C3da neurons, and found this was sufficient to induce robust axon regeneration (Figure 6C-6D). Next, we performed epistasis analysis to assess the genetic interaction between neuronal activity and Ras *in vivo*. Optogenetics-induced axon regeneration of C3da neurons was abolished after Ras^DN^ expression (Figure S7A-S7B). Moreover, parvalbumin induced inhibition of axon regeneration in C4da neurons was reversed when Ras^CA^ was co-expressed in C4da neurons (Figure S7C-S7D). These results placed Ras as a downstream effector of neuronal activity. Finally, we tested whether neuronal activity could activate Ras in larval sensory neurons. Acute application of the transient receptor potential A1 (TrpA1) channel agonist AITC, which potently increases neuronal activity and Ca^2+^ levels of C4da neurons (Gu et al., 2019; Xiang et al., 2010), induced a more than two-fold increase of dpERK in C4da neurons (Figure S7E-S7F). Together, our results indicate Ras is a neuronal type-specific injury signal that acts downstream of Ca^2+^ spikes to determine the different axon regeneration abilities of C3da and C4da neurons.

**Figure 6.**
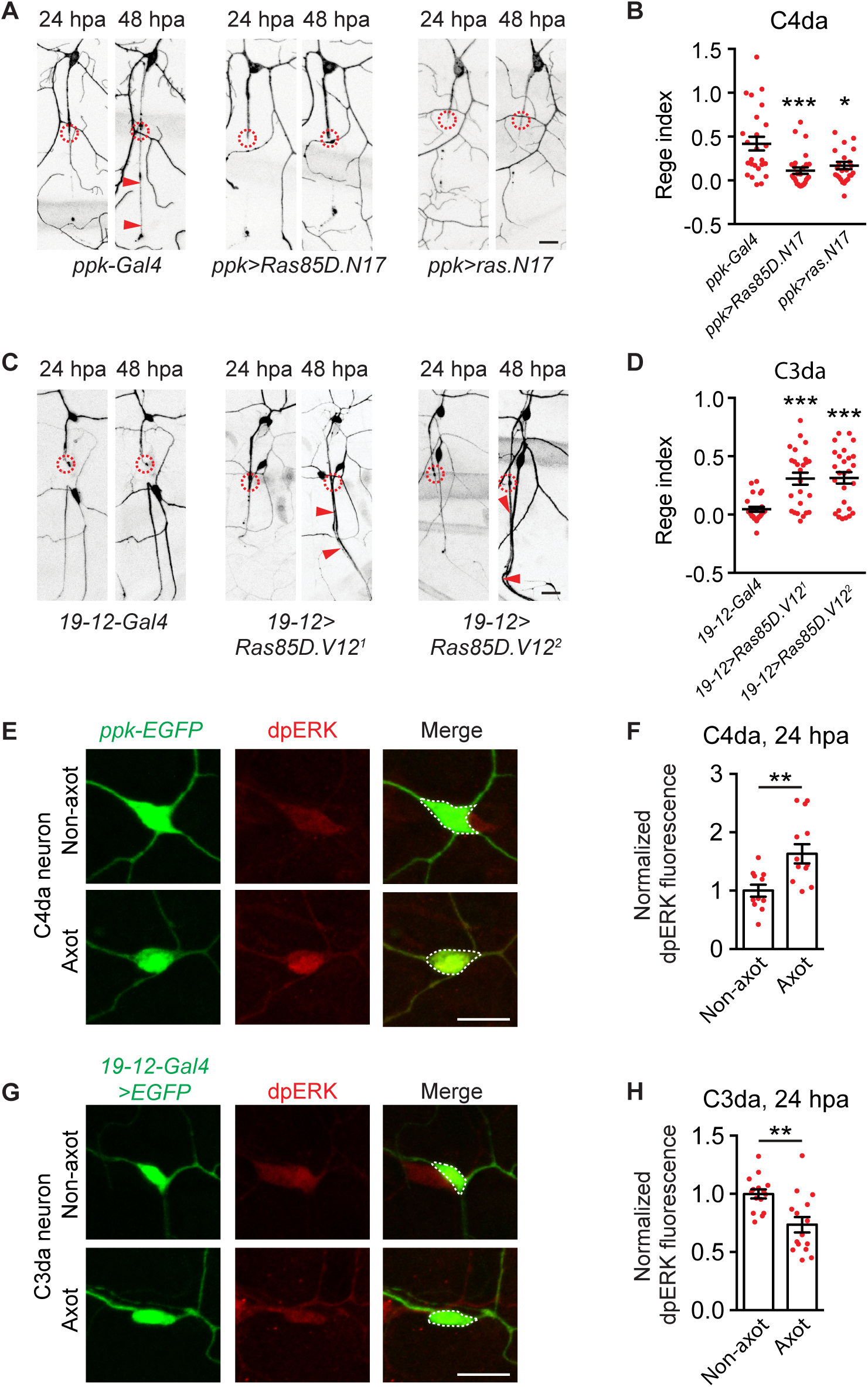
Neuronal-type specific Ras activation determines axon regeneration outcome. (A-B) Representative images (A) and regeneration index (B) of axon regeneration of C4da neurons from control larvae (*ppk-Gal4*) and larvae expressing dominant negative Ras (*ppk-Gal4>UAS-Ras85D.N17* or *ppk-Gal4>UAS-ras.N17*) in C4da neurons. Circles indicated axotomy sites and red arrowheads showed regenerated axons. n=25, 27 and 24. Scale bar, 20 µm. (C-D) Representative images (C) and regeneration index (D) of C3da neurons from control larvae (*19-12-Gal4*) and larvae expressing constitutively active Ras (*19-12-Gal4>UAS-Ras85D.V12^1^* and *19-12-Gal4>UAS-Ras85D.V12^2^*) in C3da neurons. Circles indicated axotomy sites and red arrowheads showed regenerated axons. n=26, 25 and 27. Scale bar, 20 µm. (E-F) Representative example (E) and quantification (F) of dpERK immunostaining signals from non-axotomized (control) and axotomized C4da neurons at 24 hpa. n=11 and 11. Scale bar, 10 µm. (G-H) Representative example (G) and quantification (H) of dpERK immunostaining signals from non-axotomized (control) and axotomized C3da neurons at 24 hpa. n=15 and 15. Scale bar, 10 µm. dpERK levels were quantified from somata as indicated by white dashed lines in (E) and (G). Kruskal-Wallis test followed by Dunn’s test (B and D) and Two-tailed unpaired *t*-test (F and H). * *P*<0.05, ** *P*<0.01, *** *P*<0.001, **** *P*<0.0001. n. s., not significant. Data are represented as mean ± S.E.M.. See also Figure S7.

### Adenosine receptor activation promotes RGC axon regeneration and survival in adult mouse

Adult CNS neurons in mammals fail to regenerate their axons after injury, posing a major challenge to functional recovery (He and Jin, 2016). Our *Drosophila* studies indicate that AdoR activation in C3da neurons is sufficient to convert C3da neurons from being non-regenerative to regenerative. Axon regenerative mechanisms are conserved (He and Jin, 2016), however, it is yet unknown whether activation of the mammalian ortholog of *Drosophila* AdoR could promote CNS axon regeneration. To address this question, we exploited optic nerve crush model to study RGC axon regeneration in adult mice (He and Jin, 2016). All four mammalian adenosine receptors (i.e., Adora1, Adora2a, Adora2b and Adora3) are GPCRs. Among them, Adora2a and Adora2b exhibited the highest sequence similarity to *Drosophila* AdoR (54% and 50% respectively, Figure S8). These mammalian adenosine receptors activate different intracellular signals as Adora2a and Adora2b are coupled to Gαs, whereas Adora1 and Adora3 are coupled to Gαi (Antonioli et al., 2013; Jacobson and Gao, 2006). Because *Drosophila* AdoR is proposed to function through Gαs (Dolezelova et al., 2007), our sequence and functional comparison together indicated that mammalian Adora2a and Adora2b are the closest orthologs of *Drosophila* AdoR. By analysis of the published single-cell RNA-seq dataset of mouse RGCs (Tran et al., 2019), we found that although Adora1 showed moderate expression in most RGC types, expression of Adora2a, Adora2b, and Adora3 in RGCs was sparse and was at very low levels (Figure S9). As Adora2a and Adora2b activate similar pathways, we chose Adora2b by testing whether Adora2b activation could promote RGC axon regeneration. To this goal, we ectopically expressed Adora2b in RGCs by intravitreal injection of AAV2-Adora2b virus. Fourteen days after virus injection, we crushed the optic nerve, and intravitreally injected adenosine to activate Adora2b for another two weeks (Figure 7A). We found this treatment significantly increased the number of regenerated axons at various distance from the crush site, when compared to adenosine injection or PBS injection alone groups (Figure 7B-7C). Inhibition of phosphatase and tensin homolog (PTEN), a negative regulator of mTOR, robustly promotes axon regeneration and cell survival of RGCs (Li et al., 2016; Park et al., 2008). Strikingly, we found that RGC regeneration induced by Adora2b was as strong as AAV2-PTEN-shRNA virus-induced PTEN knockdown (Figure 7B-7C). Further, combining Adora2b activation with PTEN knockdown resulted in significantly more RGCs axon regeneration when compared to either single treatment, revealing a synergistic effect (Figure 7B-7C). Optic nerve crush is followed by loss of ∼80% RGCs in two weeks (Park et al., 2008). We then determined whether Adora2b activation could protect against neuronal death. Indeed, Adora2b activation, but not adenosine injection alone, increased the number of surviving RGCs (Figure 7D-7E). Knocking down PTEN by AAV2-PTEN-shRNA virus injection also increased RGCs survival (Figure 7D-7E). Combining Adora2b activation with PTEN knockdown further increased the number of surviving RGCs, when compared to singular treatment (Figure 7D-7E), indicating a synergistic effect on cell survival. Taken together, our results indicate that activation of Adora2b signaling promotes both axon regeneration and survival of RGCs in adult mice. Because Adora2b activation in mouse RGCs and AdoR activation in *Drosophila* larval C3da neurons both promote axon regeneration, we propose a conserved role of purinergic signaling in promoting axon regeneration.

**Figure 7.**
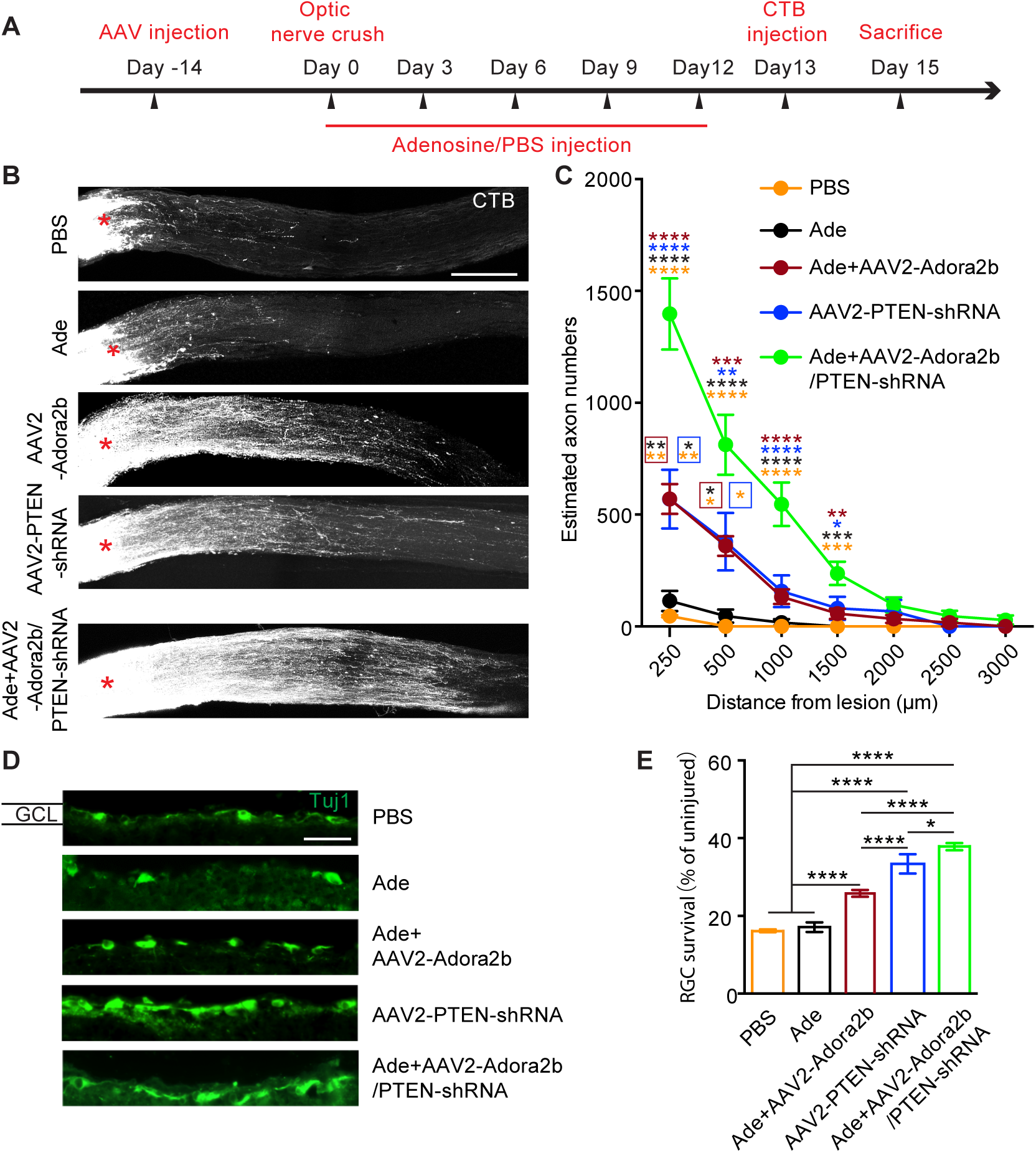
Adora2b activation promote RGC axon regeneration and survival in adult mice. (A) Schematic drawing shows the experimental procedures for studying RGC axon regeneration after optic nerve crush in adult mice. Adora2b was virally transfected, and 2 µL 10 µM adenosine (Ade) or PBS was intravitreally injected for a total of 5 times. (B) Representative confocal images of optic nerve cryosections showing regenerated RGC axons labeled by CTB, in various treatment groups. Crush site is indicated by red asterisk. Scale bar, 250 µm. (C) Quantification of RGC axon regeneration in different treatment groups. Y-axis shows the number of axons and X-axis indicates the distance distal to the crush sites. n = 9, 13, 17, 11 and 11 nerves from n=6, 8, 12, 8 and 8 mice in PBS, Ade, Ade+AAV2-Adora2b, AAV2-PTEN-shRNA, Ade+AAV2-Adora2b/PTEN-shRNA group, respectively. One-way ANOVA followed by Tukey’s test. Asterisk colors indicate the group that the *P* value was significant against (e.g., orange tested against PBS group, black tested against Ade group). (D) Representative confocal images of retinal cross sections showing surviving neurons in the ganglion cell layer (GCL) indicated by immunostaining of the neuron marker Tuj1. Scale bar, 50 µm. (E) Quantification of neuronal survival in different groups by normalizing to the neuron density observed in intact retinas. n = 83, 33, 37, 25 and 58 sections from n=10, 7, 6, 5 and 8 retinas in PBS, Ade, Ade+AAV2-Adora2b, AAV2-PTEN-shRNA, Ade+AAV2-Adora2b/PTEN-shRNA group, respectively. One-way ANOVA followed by Tukey’s test. * *P* < 0.05, \*\**P* < 0.01, *** *P* < 0.001, **** *P* < 0.0001. n. s., not significant. Data are represented as mean ± S.E.M.. See also Figure S8 and S9.

## DISCUSSION

While we now have a growing appreciation of factors that can promote axon regeneration, how the distinct regenerative abilities of neuronal types are encoded remains mysterious. Using *Drosophila* larval sensory neurons as an experimental model to investigate axon regeneration differences among neuronal types, we identified a glia-neuron interaction that differentially controls axon regeneration abilities in neurons that reside in close proximity to one another. In our working model (Figure 8), glial cells that ensheath both C4da and C3da neurons actively respond to axotomy. The resulting glial Ca^2+^ spikes lead to adenosine release (or release of ATP that is hydrolyzed to adenosine) that could potentially impact all sensory neuron types. However, the cell type-specific expression of AdoR determines that C4da neurons, but not C3da neurons, are capable of responding to gliotransmission. This neuronal type-specific responses to gliotransmission activates downstream axon regeneration programs, including AdoR activation, spontaneous burst firing and Ca^2+^ spikes, and Ras activity, in C4da neurons, but not in C3da neurons, eventually leading to axon regeneration specifically in C4da neurons. Remarkably, reactivation of these regenerative programs in regeneration-incompetent C3da neurons, by reconstitution of the C3da neuron responses to gliotransmission through ectopic AdoR expression, or by forced activation of neuronal activity or Ras, is sufficient to confer axon regeneration to C3da neurons. Finally, our studies of mouse RGC regeneration after optic nerve crush reveal a conserved role of adenosine signaling in promoting mammalian CNS axon regeneration, suggesting that targeting purinergic signaling could be a potential therapeutic strategy for CNS repair.

**Figure 8.**
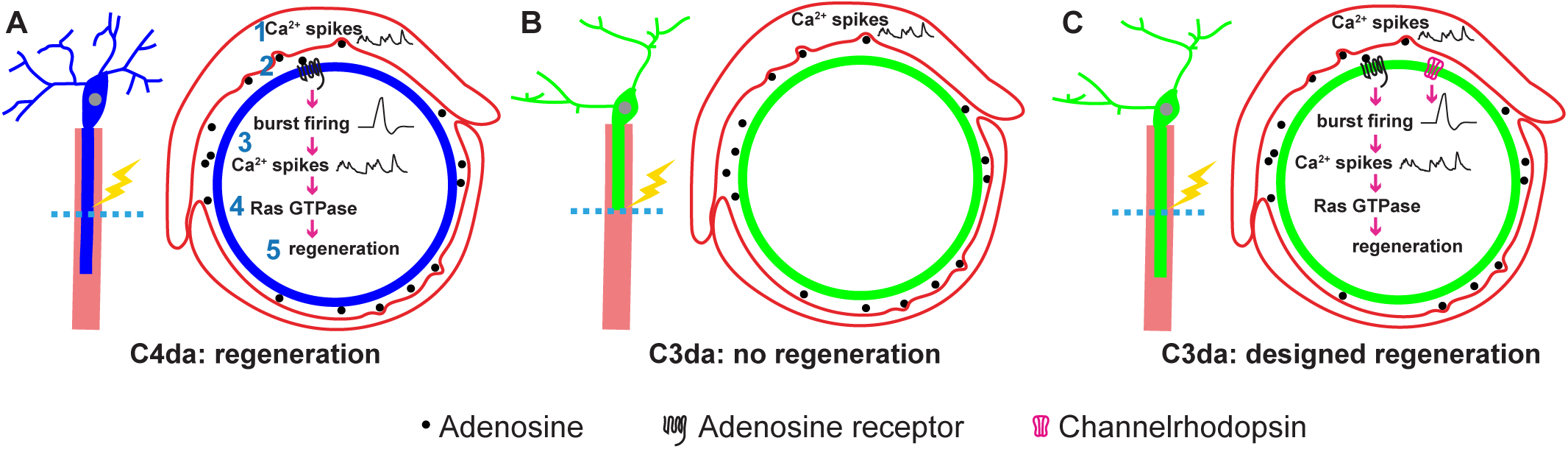
A working model for neuronal type-specific axon regeneration. (A-C) Red, blue and green colors mark ensheathing glia, C4da neurons, and C3da neurons, respectively. In (A-C), the scheme of glia-neuron organization is shown on the left panels, with laser (yellow flash) and lesion site (dashed lines) indicated. The signaling events from ensheathing glia to sensory neurons are shown in the right panels. C4da and C3da neurons differentially respond to glial chemical transmission, resulting in different regenerative responses. (A) The sequence of glia-to-C4da neuron signaling events. 1, axotomy induces glial Ca^2+^ spikes, triggering gliotransmission. 2, the gliotransmitter adenosine activates AdoR in C4da neurons. 3, leading to spontaneous neuronal activity manifested as Ca^2+^ spikes and burst firing. 4, which subsequently activates Ras, 5, through a process likely involving gene transcription, activated Ras triggers axon regeneration of C4da neurons. (B) Axotomy of C3da neurons also elicits glial Ca^2+^ spikes and gliotransmission. However, C3da neurons do not respond to gliotransmission due to lack of AdoR expression, and hence do not mount axon regenerative programs. (C) Axon regeneration of C3da neurons is induced by reconstitution of C3da responses to gliotransmission *via* ectopic AdoR expression, by optogenetic stimulation, or by genetic activation of Ras.

Both the intrinsic properties and extrinsic factors of neurons regulate axon regeneration (He and Jin, 2016). Extrinsic factors studied include glial scar, myelin-associated inhibitory factors, and glial metabolic status (Li et al., 2020; Yiu and He, 2006). As yet another important glia-neuron crosstalk, whether gliotransmission could play a role in axon regeneration is unknown. In this study, we demonstrate that gliotransmission plays an instructive role in promoting axon regeneration of select *Drosophila* neuron types, thus expanding our knowledge of how glia control axon regeneration. We also provide a precedent that neuronal types could differentially respond to gliotranmission, due to AdoR expression in selective neuronal type. Therefore, our studies reveal an elegant interplay between intrinsic properties (i.e., AdoR expression in selective neuron type) and extrinsic factors (i.e., gliotransmission) in determining whether injured neurons will regenerate its axon (Figure 8). Moreover, our studies reveal a new physiological function of gliotransmission, in addition to its well-known role in modulating synaptic strength (Bazargani and Attwell, 2016; Haydon, 2001; Lezmy et al., 2021; Ma et al., 2016; Pascual et al., 2005).

Because gliotransmission acts upstream to establish axon regenerative programs in neurons, and because gliotransmission is a result of glial Ca^2+^ signals, understanding how Ca^2+^ spikes in ensheathing glia are induced after axotomy will provide insights of how nerve injury is sensed. In both mammals and *Drosophila*, glial cells express neurotransmitter receptors (Haydon, 2001). This allows glial cells to respond to neuronal activity, leading to cytosolic Ca^2+^ rise and subsequent release of gliotransmitters (Haydon, 2001). These studies lead to the view that glia can reciprocally communicate with neurons, by ‘listening to’ neuronal activity and providing feedback regulation of neuronal functions (Haydon, 2001). Because axotomized C4da neurons exhibit burst firing, it raises a possibility that glial Ca^2+^ spikes could result from C4da neuron activity. To test this idea, we activated C4da neurons by AITC application (Gu et al., 2019; Xiang et al., 2010). However, we found that AITC did not activate ensheathing glia (Figure S2H-S2I), arguing against this possibility. Rather, our data suggest a unidirectional signaling from ensheathing glia to C4da neurons (Figure 8). We speculate that Ca^2+^ spikes in ensheathing glia could be produced *via* a cell-autonomous mechanism, or *via* a cell-non-autonomous mechanism that involves glial interactions with surrounding cells including non-neuronal cells.

*Drosophila* larval C4da and C3da neurons are functionally analogous to the polymodal nociceptors and low-threshold mechanoreceptors in mammalian DRG, respectively (Xiang et al., 2010; Yan et al., 2013). The burst firing events found in axotomized C4da neurons are reminiscent of those ectopic discharges of injured DRG neurons in the neuropathic pain model (Costigan et al., 2009). These ectopic discharges are thought to contribute to spontaneous pain, a hallmark of neuropathic pain (Costigan et al., 2009). Interestingly, gliotransmission is critically involved in the ectopic discharge of injured DRG neurons (Huang et al., 2013). Satellite glia, which wrap DRG neurons, have been shown to signal to DRG neurons through gliotransmission to modulate neuronal excitability (Huang et al., 2013). This parallel comparison between *Drosophila* and mouse suggests that burst firing of axotomized C4da neurons could reflect a nociceptive sensitization state, and this sensitization state requires gliotransmission. Therefore, a potential physiological function of gliotransmission could be to sensitize C4da nociceptors, but not C3da touch receptors, after nerve injury. This could likely explain why AdoR is expressed in C4da, but not C3da, neurons. Together, our study suggests that axon regeneration and nociceptive sensitization, two separate events following axotomy, could be mechanistically related to each other. Along this line, it would be interesting to determine whether gliotransmission, which is important for chronic pain (Huang et al., 2013), plays a role in DRG axon regeneration.

By optogenetic stimulation of C3da neurons with different firing patterns, we find that the neuronal activity pattern, instead of overall excitability increases, dictates the axon regeneration outcomes. Different patterns of neuronal activity elicit different Ca^2+^ responses. Burst firing, but not tonic firing, elicits strong Ca^2+^ spikes in C3da neurons, likely due to prolonged membrane depolarization. Our findings stress the importance of precisely patterned neuronal activity for achieving optimal axon regeneration. Stimulation of neuronal activity has been used to promote axon regeneration (Li et al., 2016; Lim et al., 2016; Zhang et al., 2019). However, the reported effects are quite variable, and the reasons of such inconsistency are not completely known (Tedeschi et al., 2016). Notably, the firing patterns in these studies are not precisely controlled, and this might contribute to the inconsistent results. The recent development of wireless optogenetics allows light to penetrate through deep tissues without an optical fiber (Montgomery et al., 2015), providing a possible means to stimulate regenerating neurons *in vivo* with a precisely controlled neuronal activity pattern. With these technological advancements, future studies are warranted to determine whether different neuronal activity patterns could exert different axon regeneration effects in other experimental models.

Axotomy-associated Ca^2+^ transients are one of the earliest neuronal injury responses (Ghosh-Roy et al., 2010; Mahar and Cavalli, 2018). The findings that C4da and C3da neurons exhibit similar immediate Ca^2+^ transients argue against a major role of immediate Ca^2+^ transients in determining axon regeneration differences among neuronal types; instead, it supports immediate Ca^2+^ transients as a general injury signal. On the other hand, Ca^2+^ spikes represent a new form of injury signal with unique characteristics. Ca^2+^ spikes exhibit neuronal type-specificity, show delayed onset but precede axon regeneration, persist throughout the axon regeneration time window, and are evoked after injury of the axon but not dendrites. Optogenetic stimulation of C3da neurons, starting at 6 hpa, separates the roles of immediate Ca^2+^ transients *versus* Ca^2+^ spikes by showing that Ca^2+^ spikes are sufficient to induce axon regeneration. Because Ca^2+^ signals of different spatiotemporal patterns can activate different signaling pathways in neurons (Berridge, 1998), we hypothesize that Ca^2+^ spikes and immediately Ca^2+^ transients regulate different aspects of axon regeneration. For example, while immediately Ca^2+^ transients could regulate cytoskeletal organization and growth cone formation (Mahar and Cavalli, 2018), persistent Ca^2+^ spikes could help maintain the Ras activity level that is required for axon regeneration of C4da neurons. It will be of interest to determine whether Ca^2+^ spikes are induced in other axon regeneration experimental models, and if so, play a role in axon regeneration.

Ras controls cell growth and is a signaling hub that can be activated by divergent signals, including receptor tyrosine kinase and G protein-coupled receptors. It is thus plausible that growth factors, neurotransmitters, and peptides can regulate axon regeneration through regulation of Ras activity. Among the many Ras targets, the canonical MAPK pathway has been shown to regulate axon regeneration, as activation of this pathway promotes axon regeneration in both mammals and flies (Hollis et al., 2009; O’Donovan et al., 2014; Perlson et al., 2005; Wang et al., 2020), and activated ERK has been found to transport retrogradely from the injury site to the cell body (Perlson et al., 2005).

Adenosine signaling has been proposed to exert protective functions in diverse tissues including the nervous system (Antonioli et al., 2013), by acting through its receptors to trigger diverse signaling events. The adenosine concentration in the extracellular space is normally low but can be rapidly increased after tissue injury (Antonioli et al., 2013). A previous study has shown that pharmacological activation of Adora3, but not Adora1, Adora2a or Adora2b, promotes neurite outgrowth of cultured RGCs (Nakashima et al., 2018). In the current study, we find that Adora2b activation promotes axon regeneration of mouse RGCs and protects against neuronal death. Importantly, the effect of Adora2b activation on axon regeneration is comparable to that after PTEN knockdown, a treatment that induces reliable and robust CNS axon regeneration by elevating mTOR activity (He and Jin, 2016). Moreover, combining Adora2b activation with PTEN knockdown further enhanced RGC axon regeneration. These findings indicate that Adora2b activation, either alone or in combination with other pathways, could be a promising strategy for mammalian CNS repair.

Neuronal type-specific regeneration is broadly observed, occurring in both CNS and PNS and in both vertebrates and invertebrates. Our studies reveal gliotransmission as novel glia-neuron crosstalk that specifies different axon regenerative abilities among neuron types. Because gliotransmission is a conserved phenomenon, future studies are needed to determine whether gliotransmission could regulate axon regeneration in other experimental systems and in other species, and whether gliotransmitters other than ATP/adenosine could be involved.

## ACKNOWLEDGEMENTS

Authors are grateful to Dr. B. Ye for help with *in vivo* optogenetic stimulation, Drs. M. Zurovec, N. Perrimon, C. Collins, C. Wu and A. Bergmann for reagents; Q. Wen and G. Dai for help on Matlab coding; Y. Peng for RNAseq data analysis. We would also like to thank A. Byrne, M. Francis, B. Ye, W. Ge, and K. Thompson-Peer for critical comments on the manuscript; Y. Song for advice on laser axotomy; Bloomington and Vienna *Drosophila* Resource Center for fly stocks. Y.L. is supported by Shanghai Municipal Science and Technology Major Project No. 2018SHZDZX05. Y.X. is supported by the National Institutes of Health under award number 1R21NS107924, the Human Frontier Science Program Young Investigator award RGY0090/2014, and the Martina Stern Memorial Fund.

## AUTHOR CONTRIBUTIONS

Conceptualization, F.W., K.T.P., and Y.X.; Methodology, F.W., K.T.P., S.Z., and Y.Q.; Investigation, F.W., K.T.P., S.Z., Y.Q., Y.L., and Y.X.; Formal Analysis, F.W., K.T.P., S.Z., Y.Q., Y.L., and Y.X.; Resources, J.G. and Y.S.; Writing—Original Draft, F.W., K.T.P., and Y.X.; Writing—Review & Editing, F.W., K.T.P., S.Z., Y.L., and Y.X.; Supervision, Y. L. and Y.X.; Funding Acquisition, Y.L. and Y.X.

## DECLARATION OF INTERESTS

The authors declare no conflicts of interest.

---

**Figure S1.**
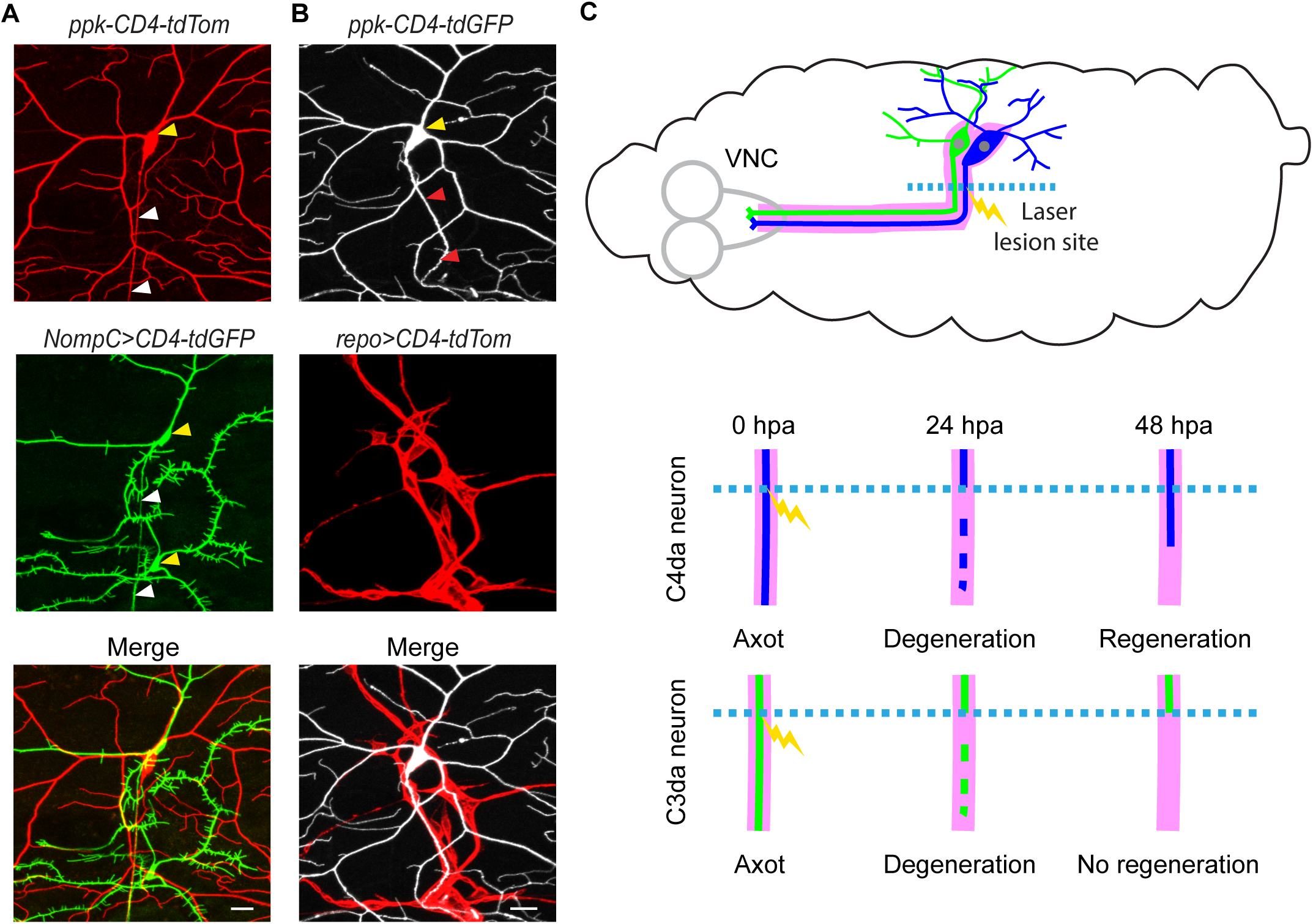
The axotomy diagram, related to Figure 1. (A) Representative images showing C4da and C3da neurons from the dorsal cluster of larval PNS. One C4da (i.e., ddaC, top panel) and two C3da neurons (upper, ddaF; lower, ddaA, middle panel) were shown, with somata close to each other and dendrites overlapped. White and yellow arrowheads indicated the axon and cell body, respectively. ddaC and ddaF were used in the current study. *ppk-CD4-tdTomato* and *NompC-Gal4>CD4-tdGFP* mark C4da and C3da neurons, respectively. Scale bar, 20 µm. (B) Representative images showing glial ensheathment of larval PNS sensory neurons. C4da neuron and ensheathing glia were marked by *ppk-CD4-tdGFP* and *repo-Gal4>UAS-CD4-tdTomato*, respectively. Red and yellow arrowheads indicated the axon and cell body, respectively. Scale bar, 20 µm. (C) Upper, a schematic diagram showing glia (magenta) ensheath axons of sensory neurons (green and blue). VNC, ventral nerve cord. Laser (yellow flash) was delivered for *in vivo* axotomy. Lower, the distal axon degenerated by 24 hpa, and regrowth of the proximal axon between 24 and 48 hpa was quantified. For clarity, only a single axon was shown.

**Figure S2.**
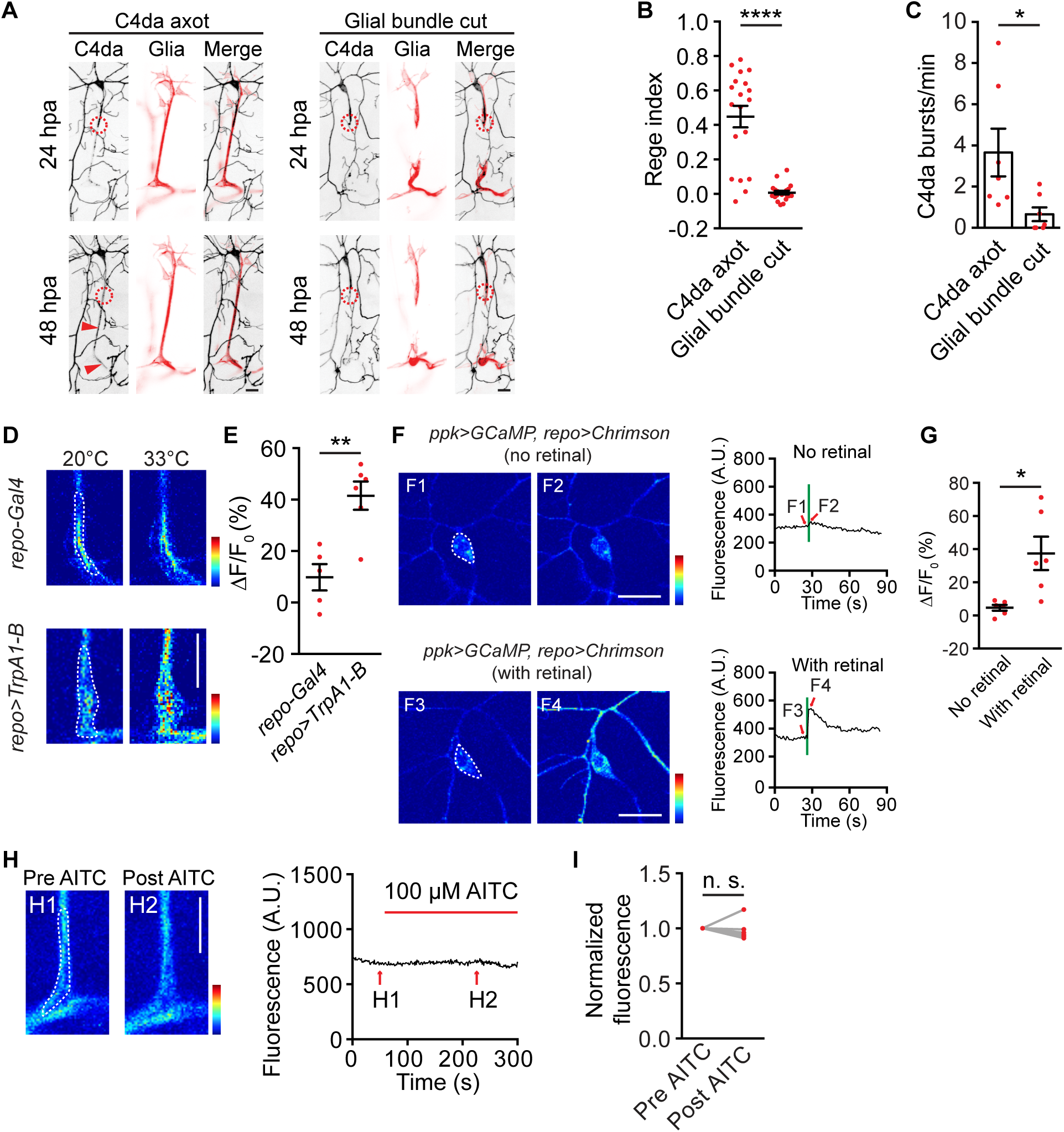
Glia-neuron crosstalk, related to Figure 1 and 2. (A) Representative images of axon regeneration of C4da neurons as well as morphology of ensheathing glia after standard laser axotomy of C4da neurons (C4da axot), and after treatment with increased laser area that also severed ensheathing glia (Glial bundle cut). In the ‘Glial bundle cut’ scenario, ensheathing glia were damaged and C4da axon regeneration was completely abolished. Circles indicated laser injury sites and red arrowheads showed regenerated axons. Scale bar, 20 µm. (B) Quantification of regeneration index of C4da neurons in ‘C4da axot’ and ‘Glial bundle cut’ modes. n=19 and 21. Mann-Whitney test. (C) Quantification of the frequency of spontaneous burst firing of axotomized C4da neurons measured by extracellular recording in ‘C4da axotomy’ and ‘Glial bundle cut’ modes. n=7 and 7. Mann-Whitney test. (D) Representative glial Ca^2+^ signals in control larvae (*repo-Gal4*) and larvae expressing TrpA1-B in glia (*repo-Gal4>UAS-TrpA1-B*), in response to thermogenetic stimulation. Scale bar, 20 µm. Color scale indicated dynamic range from 50 to 1,600. Regions of interest were indicated by white dashed lines. (E) Quantification of glial Ca^2+^ responses to thermogenetic stimulation. n=5 and 6. Two-tailed unpaired *t*-test. (F-G) Representative images (F) and summary (G) of Ca^2+^ signals of C4da neurons in response to acute optogenetic stimulation of CsChrimson-expressing glia. Green light (5.0 mW/mm^2^, 555 nm light pattern with 500 ms on and 500 ms off was repeated 10 times) was delivered. Larvae were fed with either no retinal (control) or 400 µM retinal (optogenetics group). Summary results in (G) shows peak responses quantified from somata of C4da neurons as indicated by dashed lines in (F). Scale bar, 20 µm. Color scale indicated dynamic range from 0 to 2,000. n=6 and 6. Two-tailed unpaired *t*-test. (H-I) Representative images (H) and summary (I) of glial Ca^2+^ responses to 100 µM AITC application. H1 and H2 show glial Ca^2+^ signals before and during AITC application, respectively. Scale bar, 20 µm. Color scale indicated dynamic range from 0 to 3,000. Regions of interest were indicated by white dashed lines. n=6. Wilcoxon matched-pairs signed rank test. * *P*<0.05, ** *P*<0.01, *** *P*<0.001, **** *P*<0.0001. n. s., not significant. Data are represented as mean ± S.E.M..

**Figure S3.**
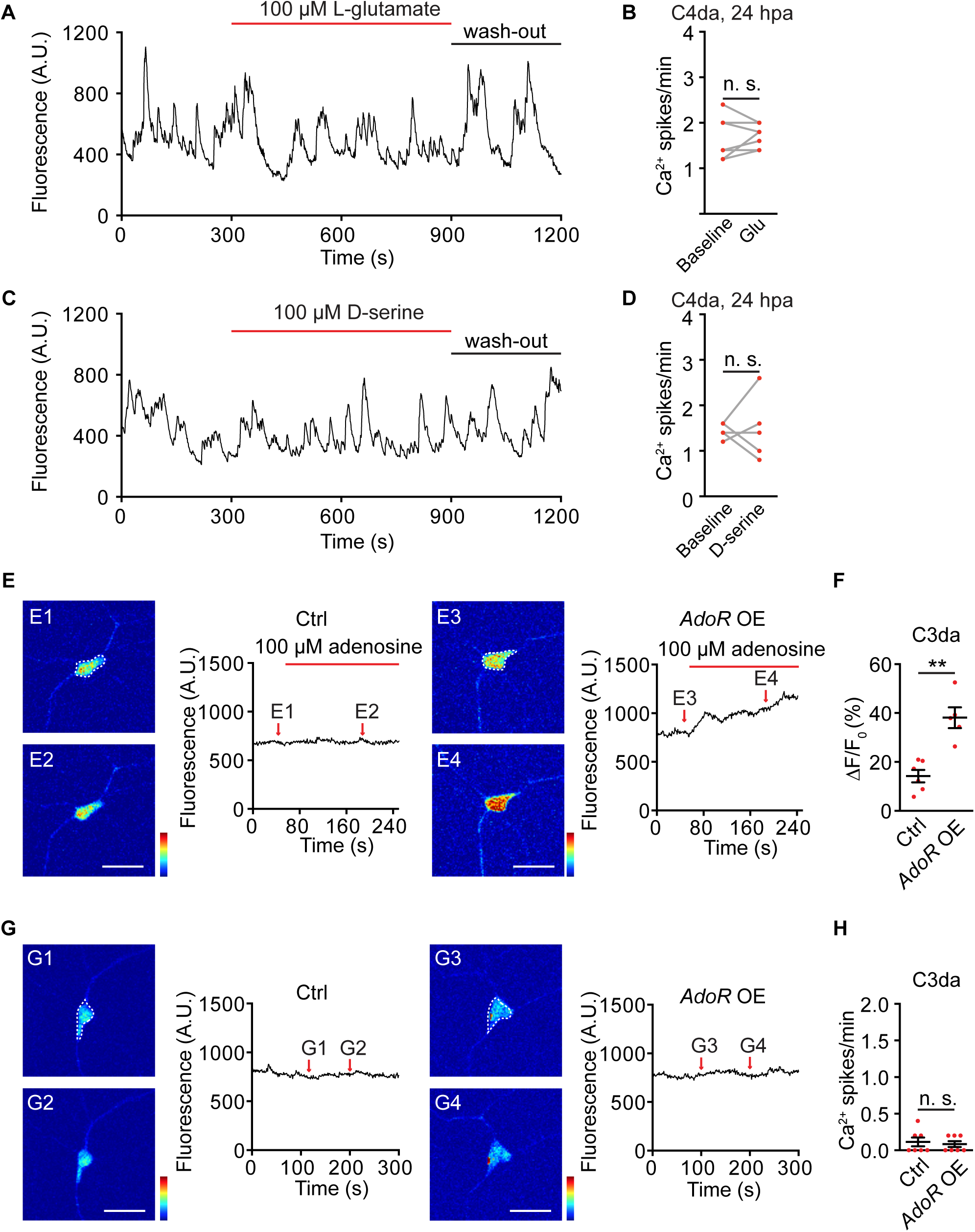
Neuronal responses to gliotransmitters, related to Figure 2 and 4. (A-D) Representative time-lapse GCaMP6s traces and summary results of Ca^2+^ responses of axotomized C4da neurons to 100 µM L-glutamate (A-B) and 100 µM D-serine (C-D) application. The Ca^2+^ spike frequency of axotomized C4da neurons was quantified from the somata before and during drug application. n=7 (B) and n=6 (D). Two-tailed paired *t*-test. (E-F) Representative Ca^2+^ signals (E) and summary (F) of non-axotomized C3da neurons to 100 µM adenosine application, from wildtype larvae (control, *19-12-Gal4>UAS-GCaMP6s*) and larvae overexpressing AdoR (AdoR OE, *19-12-Gal4>UAS-AdoR, GCaMP6s*) in C3da neurons. E1-E4 show individual image frames derived from the associated time-lapse GCaMP6s imaging traces shown in the right. Data were quantified from the somata of C3da neurons indicated by white dashed lines. Color scale indicated dynamic range from 50 to 1,700. n=6 and 5. Two-tailed unpaired *t*-test. Scale bar, 20 µm. (G-H) Representative (G) and summary (H) of spontaneous Ca^2+^ signals of non-axotomized C3da neurons, from wildtype larvae (control, *19-12-Gal4>UAS-GCaMP6s*) and larvae overexpressing AdoR (AdoR OE, *19-12-Gal4>UAS-AdoR, GCaMP6s*) in C3da neurons. G1-G4 show individual image frames derived from the associated time-lapse GCaMP6s traces shown in the right. Data were quantified from the somata of C3da neurons as indicated by dashed lines. Color scale indicated dynamic range from 0 to 3,000. n=7 and 7. Mann-Whitney test. Scale bar, 20 µm. * *P*<0.05, ** *P*<0.01, *** *P*<0.001, **** *P*<0.0001. n. s., not significant. Data are represented as mean ± S.E.M..

**Figure S4.**
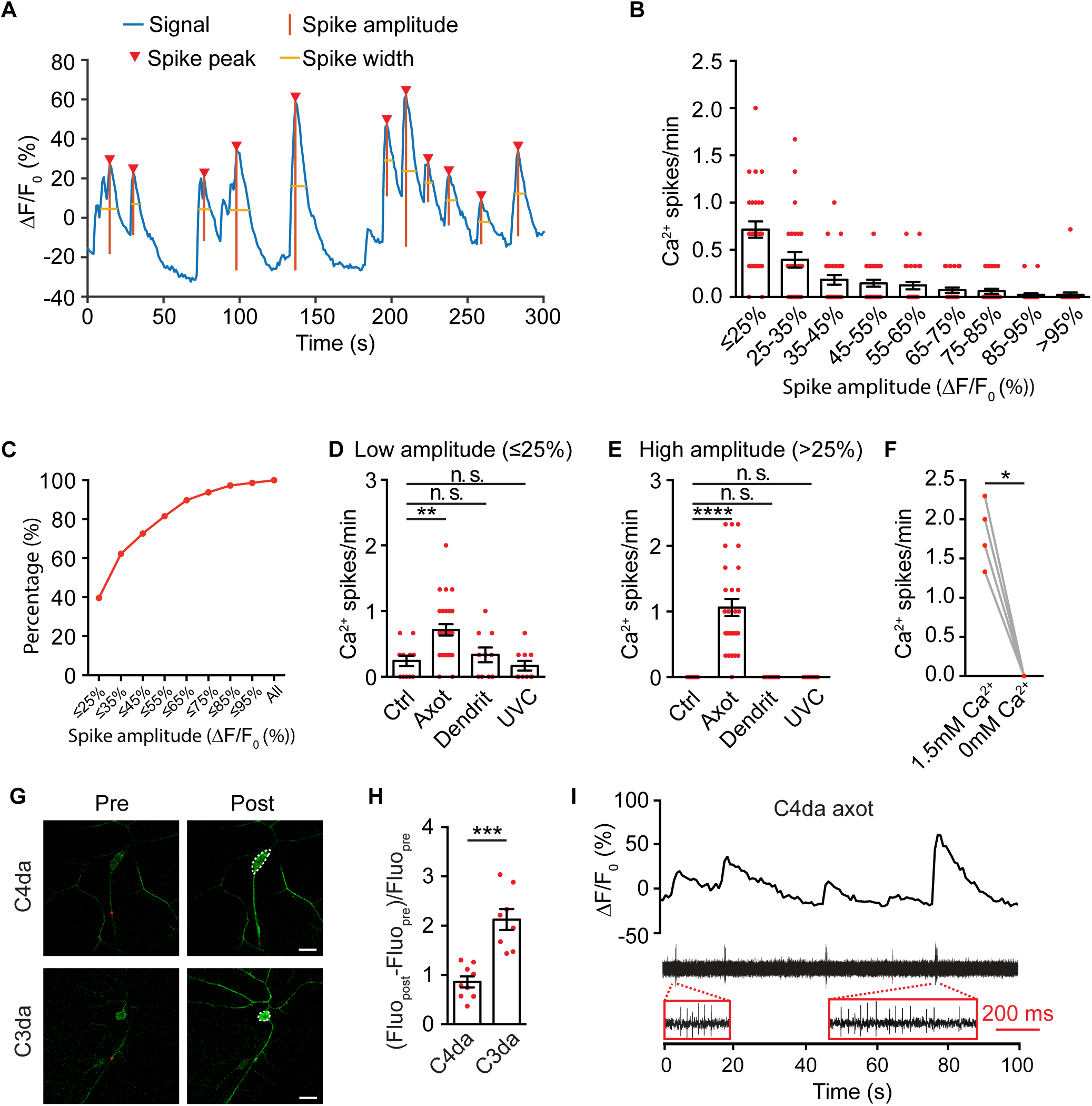
The characteristics of Ca^2+^ spikes, related to Figure 3. (A) Ca^2+^ spike detection. A representative somatic GCaMP6s traces of axotomized C4da neurons, with Ca^2+^ spikes being detected by the ‘Findpeaks’ function of Matlab. Blue curve showed the normalized GCaMP6s signal. Red triangles marked the peak position of individual Ca^2+^ spikes. Red lines (vertical) and orange lines (horizontal) indicate spike amplitude (prominence) and spike width, respectively. See Video S3, S4 and Star Methods for details. (B) Distribution of the average frequency of Ca^2+^ spikes (Y axis) over the spike amplitude (X axis), of axotomized C4da neurons at 24 hpa. n=27. (C) Accumulative distribution of the average frequency of Ca^2+^ spikes (Y axis) in relationship to the spike amplitude (X axis), of C4da neuron at 24 hpa. n=146. For example, ΔF/F_0_ ≤25% referred to the amplitude of Ca^2+^ spikes no more than 25%. The percentage of all spikes falling into this group was 39.7%. (D-E) Results of Ca^2+^ spikes of axotomized C4da neurons in Figure 2E were divided into low and high amplitude groups with a cutoff of ΔF/F_0_=25%. Only the C4da axotomy group showed Ca^2+^ spikes with high amplitude. n=11, 27, 10 and 10 for each treatment group. One-way ANOVA followed by Bonferroni’s tests between non-axotomized (control) group and other groups. (F) Removing external Ca^2+^ from 1.5 mM to 0 mM blocked Ca^2+^ spikes in axotomized C4da neurons at 24 hpa. n=7. Wilcoxon matched-pairs signed rank test. (G-H) Representative images (G) and quantification (H) of peak responses of immediate Ca^2+^ transients following axotomy of C4da and C3da neurons. n=9 and 8. Two-tailed unpaired *t*-test. Scale bar, 20 µm. (I) Simultaneous GCaMP6s imaging and extracellular recording on the same axotomized C4da neurons at 24 hpa showed close temporal correlation between Ca^2+^ spikes and burst firing events. GCaMP6s signals obtained from the soma were shown in the upper panel, and electrophysiological recording traces were shown in the lower panel. Zoom-in (red box) views showed each Ca^2+^ spike was associated with a burst firing event. Scale bar in red boxes, 200 ms. See also Video S5. * *P*<0.05, ** *P*<0.01, *** *P*<0.001, **** *P*<0.0001. Data are represented as mean ± S.E.M..

**Figure S5.**
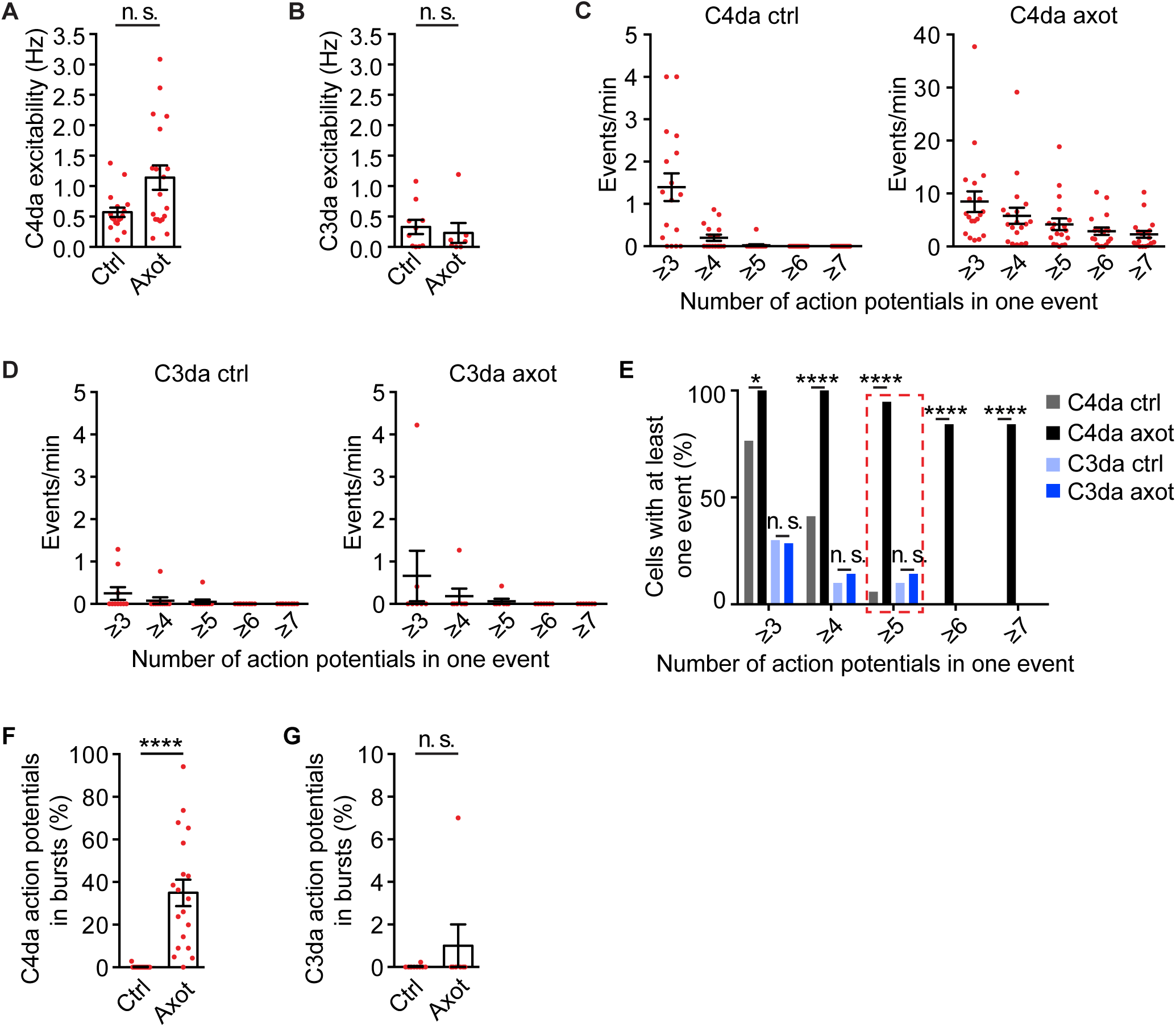
Axotomy elicited burst firing in C4da, but not C3da, neurons, related to Figure 3. (A) The excitability of C4da neurons, indicated by the overall firing frequency. n=17 for control C4da neurons and n=19 for axotomized C4da neurons. Mann-Whitney test. (B) The excitability of C3da neurons, indicated by the overall firing frequency. n=10 for control C3da neurons and n=7 for axotomized C3da neurons. Mann-Whitney test. (C-D) Average number of events per minute with indicated number of action potentials (‘≥3’ means 3 or more, others follow the same rule), recorded from control (non-axotomized) and axotomized C4da (C) and C3da neurons (D). n=17 for control and n=19 for axotomized C4da neurons; n=10 for control and n=7 for axotomized C3da neurons. Here, we define a series of consecutive action potentials (3 or more) as an “event”, on condition that the inter-spike interval of each two adjacent action potentials is less than 100 ms. (E) The percentage of neurons showing at least one event with indicated action potential number. Events with five action potentials or more were defined as a ‘burst firing event’ in this study. By this definition, burst firing was seldom detected in control C4da neurons (1 out of 17), but the possibility was significantly increased in axotomized C4da neurons (18 out of 19). Events with six action potentials or more were only detected in axotomized C4da neurons, but not in control C4da neurons, or in control or axotomized C3da neurons. n=17 for control and n=19 for axotomized C4da neurons; n=10 for control and n=7 for axotomized C3da neurons. Fisher’s exact test. (F) Percentage of action potentials that was found in the burst firing events in control (0.17%) and axotomized (35.0%) C4da neurons. n=17 and 19. Mann-Whitney test. (G) Percentage of action potentials that was found in the burst firing events in control (0.023%) and axotomized (1.0%) C3da neurons. n=10 and 7. Mann-Whitney test. * *P*<0.05, ** *P*<0.01, *** *P*<0.001, **** *P*<0.0001. n. s., not significant. Data are represented as mean ± S.E.M..

**Figure S6.**
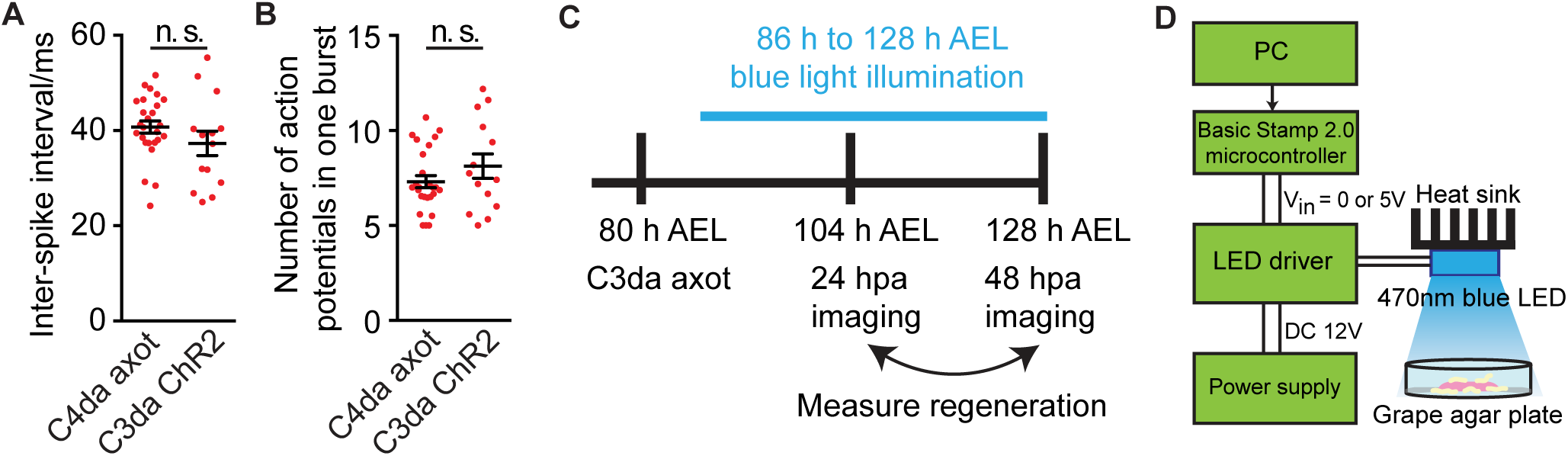
Optogenetic Stimulation Setup, related to Figure 5. (A-B) A specific blue light stimulation pattern (300 ms pulse, 470 nm light at 1.4 mW/mm^2^) triggered burst firing in C3da neurons, with inter-spike interval (A) and number of action potentials within a burst (B) resembling those recorded from axotomized C4da neurons at 24 hpa. n=26 and 14. Two-tailed unpaired *t*-test. Data are represented as mean ± S.E.M.. n. s., not significant. (C-D) A schematic experimental procedure (C) and setup (D) for *in vivo* optogenetic stimulation of free moving larvae.

**Figure S7.**
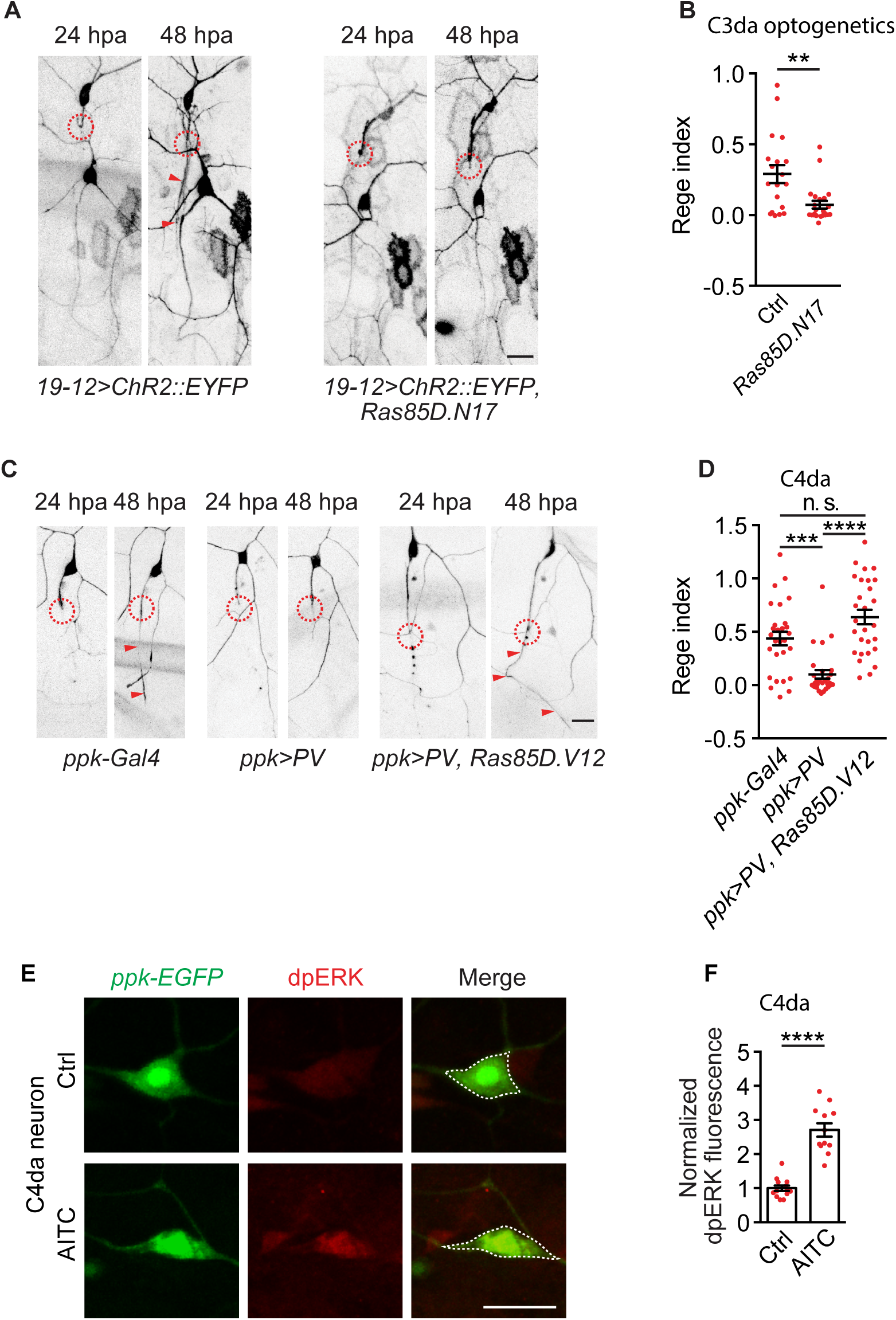
Ras acts downstream of Ca^2+^ signals to mediate axon regeneration, related to Figure 6. (A-B) Representative images (A) and regeneration index (B) of optogenetics-induced axon regeneration of C3da neurons with or without expression of dominant negative Ras in C3da neurons. Circles indicated axotomy sites and red arrowheads showed regenerated axons, same as below. 1.4 mW/mm^2^ 300 ms duration blue light was delivered to free moving larvae every 15 s. n=19 and 23. Mann-Whitney test. Scale bar, 20 µm. (C-D) Representative images (C) and regeneration index (D) of C4da neurons from control larvae (*ppk-Gal4*), larvae expressing parvalbumin in C4da neurons, and larvae expressing both parvalbumin and constitutive active Ras in C4da neurons. n=28, 28 and 28. Kruskal-Wallis test followed by Dunn’s test. Scale bar, 20 µm. (E-F) Representative examples (E) and quantification (F) of dpERK immunostaining signals in C4da neurons in response to 100 µM AITC incubation for 15 minutes. n=13 and 12. Mann-Whitney test. Scale bar, 10 µm. \**P*<0.05, \*\**P*<0.01, *** *P*<0.001, **** *P*<0.0001. n. s., not significant. Data are represented as mean ± S.E.M..

**Figure S8.**
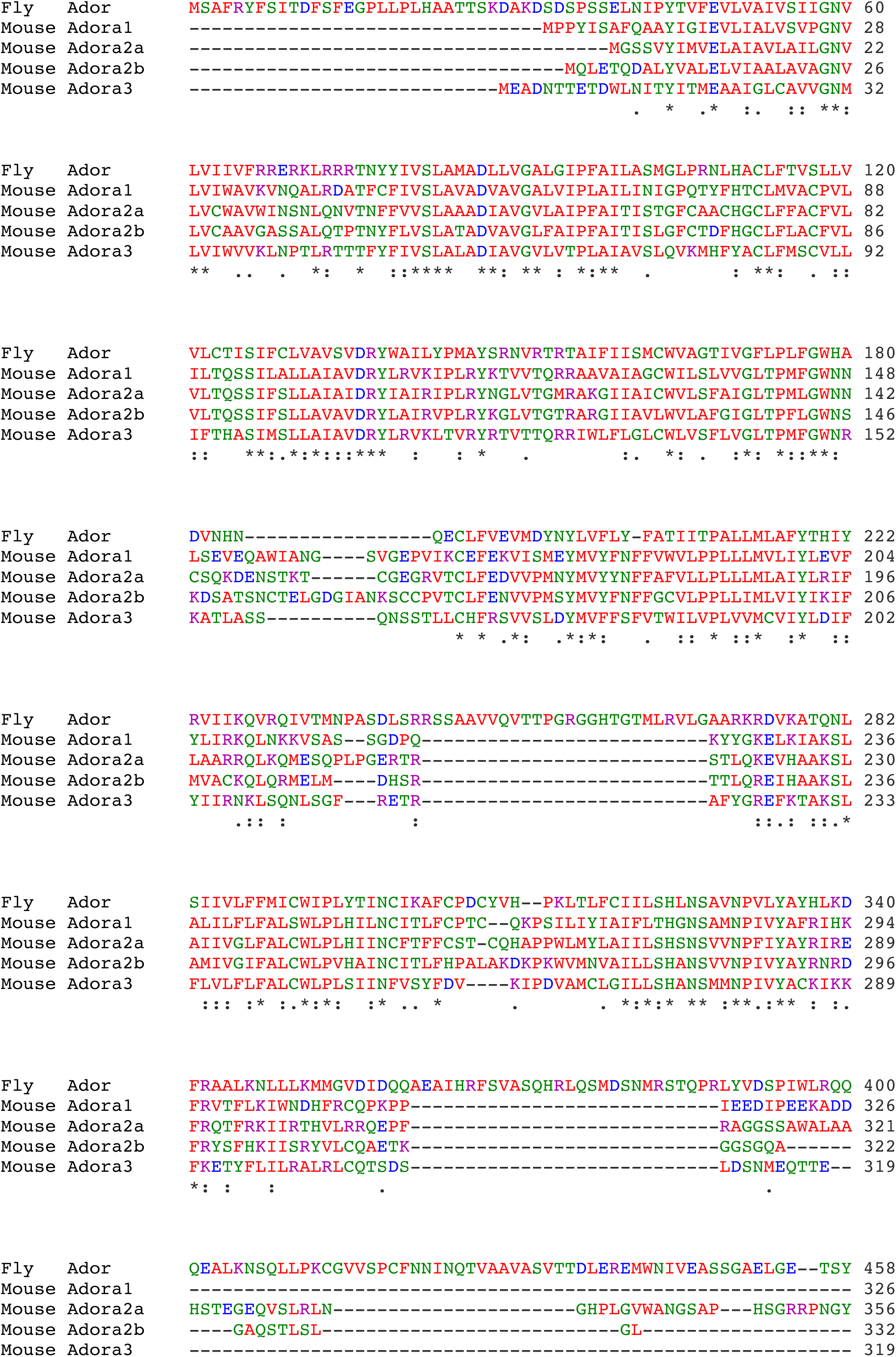
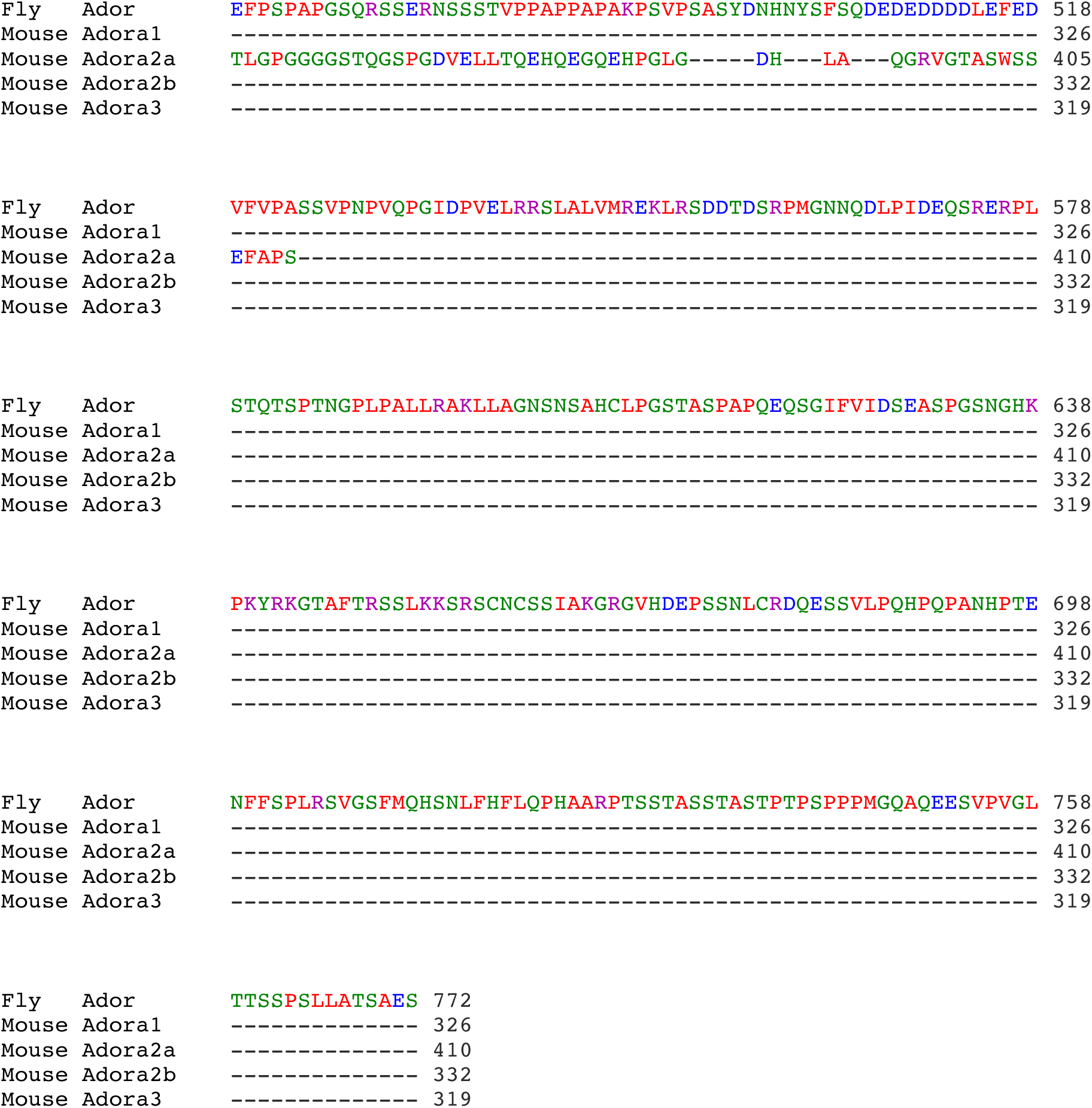
Sequence alignment of mouse adenosine receptors and *Drosophila* AdoR, related to Figure 7. The N-terminal part of the *Drosophila* AdoR protein (around 300 amino acids) comprises the region with most conservation among species and contains the seven transmembrane helices. Unlike other known adenosine receptors, *Drosophila* AdoR has a long (predicted intracellular) C-terminal extension of unknown function (Dolezelova et al., 2007). The alignment is based on previous published toolkit (Madeira et al., 2019) . Color reference means the following in amino acid residues: red, small and hydrophobic (including AVFPMILW) amino acids; blue, acidic amino acids (including DE); magenta, basic amino acids (including RK); green, Hydroxyl+sulfhydryl+amine amino acids (including STYHCNGQ). An asterisk (**\***) indicates positions, which have a single, fully conserved residue. A colon (:) indicates conservation between groups with strongly similar properties as below - roughly equivalent to scoring >0.5 in the Gonnet PAM 250 matrix. A period (.) indicates conservation between groups of weakly similar properties as below - roughly equivalent to scoring ≤0.5 and >0 in the Gonnet PAM 250 matrix.

**Figure S9.**
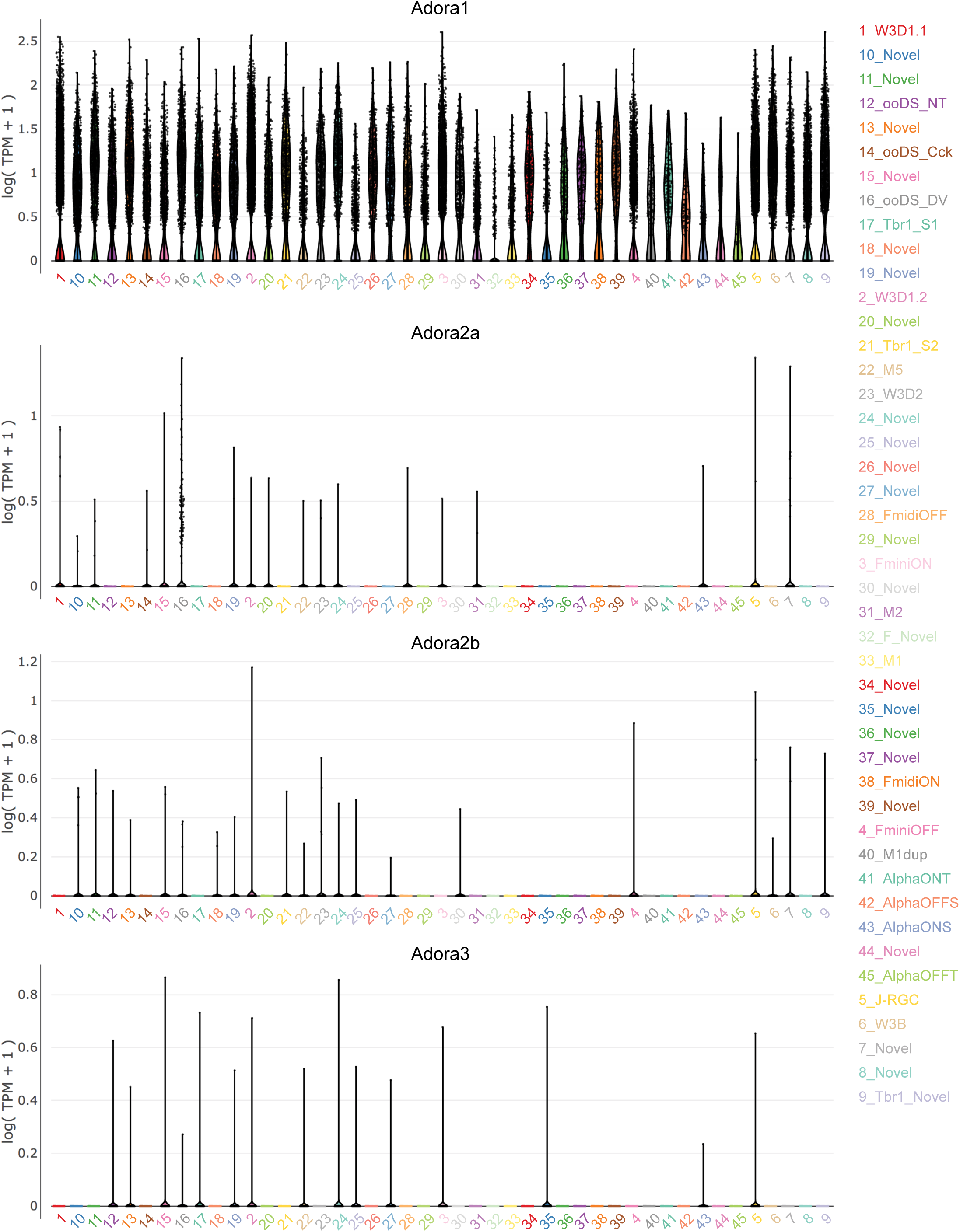
Expression of adenosine receptors in adult mouse RGCs by single-cell RNAseq, related to Figure 7. Previous report identified 45 clusters of RGC types from postnatal day 56 mice based on single-cell RNAseq, named as C1-C45 (Tran et al., 2019). The y axis is the expression level of the gene of interest with Transcripts per Million (TPM). The x axis labels all 45 clusters with distinct and corresponding colors. Individual dots show the expression level of individual cells. The cumulative distribution of expression is outlined with a violin shape. In contrast to Adora1, which is found in most RGC types, the expressions of Adora2a, Adora2b and Adora3 are sparse and very low. Figures and data are derived from: https://singlecell.broadinstitute.org/single_cell/study/SCP509/mouse-retinal-ganglion-cell-adult-atlas-and-optic-nerve-crush-time-series#study-visualize

## STAR METHODS

### KEY RESOURCE TABLE

| REAGENT or RESOURCE | SOURCE | IDENTIFIER |
| --- | --- | --- |
| Antibodies |  |  |
| Rabbit anti-Phospho-p44/42 MAPK (Erk1/2) (Thr202/Tyr204) | Cell Signaling | #4370, RRID:AB_2315112 |
| Goat anti-rabbit Alexa Fluor Plus 647 | Jackson Immunoresearch | #111-605-144, RRID:AB_2338078 |
| Mouse anti-Tuj1 (tubulin beta 3) | BioLegend | #801202, RRID:AB_10063408 |
| Donkey anti-mouse Alexa Fluor Plus 488 | Abcam | #150109 |
| Chemicals, Peptides, and Recombinant Proteins |  |  |
| Paraformaldehyde | Sigma | P6148 |
| Mounting media | Thermo Fisher | P36961 |
| Triton X-100 | Fluka | BP151 |
| Normal donkey serum | Sigma | D9663 |
| Phosphate buffered saline (powder) | Sigma | P3813 |
| Bovine serum albumin | Sigma | A2058 |
| All-trans retinal | Sigma | R2500 |
| Agarose | Thermo Fisher | BP160-500 |
| Sevoflurane | Patterson Veterinary | NDC-14043-110-25 |
| Adenosine | Sigma | A9251 |
| L-glutamate | Sigma | G1251 |
| D-serine | Sigma | S4250 |
| Cholera Toxin Subunit B, Alexa Fluor 555 Conjugate (CTB-555) | Thermo Fisher | C34776 |
| Tetrahydrofuran | Sigma | 186562 |
| Dichloromethane | Sigma | 270997 |
| DiBenzy Ether | Sigma | 108014 |
| Optimal cutting temperature compound | Sakura Finetek | 4583 |
| Bacterial and virus strains |  |  |
| rAAV2-hSyn-PTEN-shRNA | (Zukor et al., 2013) | N/A |
| rAAV2-hSyn-Adora2b | Virus core at Center for Excellence in Brain Science and Intelligence Technology, Chinese Academy of Science, Shanghai | This paper |
| Experimental Models: Organisms/Strains |  |  |
| *Drosophila*, *ppk-Gal4* | (Han et al., 2012) | N/A |
| *Drosophila*, *ppk-LexA* | (Gou et al., 2014) | N/A |
| *Drosophila*, *NompC-Gal4* | Bloomington<br>Drosophila Stock<br>Center | #BL36361 |
| *Drosophila*, *ppk-CD4-tdGFP* | (Han et al., 2011) | N/A |
| *Drosophila*, *ppk-CD4-tdTomato* | (Han et al., 2011) | N/A |
| *Drosophila*, *UAS-CD4-tdGFP* | (Han et al., 2011) | N/A |
| *Drosophila*, *UAS-CD4-tdTomato* | (Han et al., 2011) | N/A |
| *Drosophila*, *UAS-mCD8-GFP* | (Xiang et al., 2010) | N/A |
| *Drosophila*, *ppk-EGFP* | (Grueber et al., 2003) | N/A |
| *Drosophila*, *LexAop2-myr-GCaMP6s* | Laboratory of Marc<br>Freeman | N/A |
| *Drosophila*, *19-12-Gal4* | (Xiang et al., 2010) | N/A |
| *Drosophila*, *Repo-Gal80* | (Awasaki et al.,<br>2008) | N/A |
| *Drosophila*, *Repo-Gal4* | Bloomington<br>Drosophila Stock<br>Center | #B7415 |
| *Drosophila*, *NompC-LexA* | Bloomington<br>Drosophila Stock<br>Center | #BL52241 |
| *Drosophila*, *UAS-2×EYFP* | Bloomington<br>Drosophila Stock<br>Center | #BL6659 |
| *Drosophila*, *UAS-2×EGFP* | Bloomington<br>Drosophila Stock<br>Center | #BL5430, #BL5431 |
| *Drosophila*, *UAS-GCaMP6s* | Bloomington<br>Drosophila Stock<br>Center | #BL42746, #BL<br>42749 |
| *Drosophila*, *UAS-TrpA1-B* | (Gu et al., 2019) | N/A |
| *Drosophila*, *UAS-AdoR* | (Dolezelova et al.,<br>2007) | N/A |
| *Drosophila*, *UAS-Kir2.1* | (Yang et al., 2014) | N/A |
| *Drosophila*, *UAS-ChR2::EYFP* | (Hwang et al., 2007) | N/A |
| *Drosophila*, *UAS-ChR2* | Bloomington<br>Drosophila Stock<br>Center | #BL52258 |
| *Drosophila*, *UAS-CsChrimson:mCherry* | Bloomington<br>Drosophila Stock<br>Center | #BL82181 |
| *Drosophila*, *UAS-Parvalbumin* | Bloomington<br>Drosophila Stock<br>Center | #BL25030 |
| *Drosophila*, *UAS-Ras85D.N17* | Bloomington<br>Drosophila Stock<br>Center | #BL4845 |
| *Drosophila*, UAS-ras.N17 | Bloomington<br>Drosophila Stock<br>Center | #BL4846 |
| *Drosophila*, UAS-Ras85D.V12 | Bloomington<br>Drosophila Stock<br>Center | #BL4847, #BL64195 |
| *Drosophila*, UAS-TNT | Bloomington<br>Drosophila Stock<br>Center | #BL28838 |
| *Drosophila*, AdoR-t2A-Gal4 | Bloomington<br>Drosophila Stock<br>Center | #BL86138 |
| *Drosophila*, AdoR (*attP* deletion mutant) | Bloomington<br>Drosophila Stock<br>Center | #BL84447 |
| Mouse: C57BL/6J | Shanghai Laboratory<br>Animal Center,<br>Chinese Academy of<br>Sciences | N/A |
| Software and Algorithms |  |  |
| MATLAB | Mathworks | https://www.mathworks.com/ |
| pClamp 10 | Molecular Devices | https://www.moleculardevices.com/ |
| GraphPad Prism 9 | GraphPad | https://www.graphpad.com |
| Zen | Zeiss | N/A |
| ImageJ | https://imagej.nih.gov/ij/ | N/A |
| BASIC Stamp editor v2.5 | www.parallax.com | N/A |
| Adobe Illustrator | www.adobe.com | N/A |
| Adobe Premiere | www.adobe.com | N/A |
| Other |  |  |
| *Drosophila* low yeast brown food | Drosophila Resource<br>Facility at<br>UMassMed | https://www.umassmed.edu/research/cores/drosophila-resource-facility/media-recipes/ |
| DigiTherm 38-liter Heating/Cooling Incubator (for optogenetic experiment) | Tritech research | DT2-MP-47L |
| Dissection scope | Zeiss | Discovery.V8 |
| UVC crosslinker | Spectroline | Spectorlinker, 254 nm, XL-1000 |
| Hand-held UV spectrophotometer | Spectroline | AccuMAX XS-254 |
| Two-photon microscope | Zeiss | LSM 7 MP |
| Confocal fluorescence microscope | Zeiss | LSM 700 |
| Fluorescence microscope | Zeiss | Axio Examiner.D1 |
| Objective W Plan-Apochromat 20x/1.0 DIC<br>M27 75mm parfocal length 75mm<br>(FWD=1.8mm) | Zeiss | 421452-9800 |
| Objective W N-Achroplan 40x/0.75 M27<br>(FWD=2.1mm) | Zeiss | 420967-9900 |
| Compact light source (Mercury lamp) | Zeiss | HXP-120 V |
| Microelectrode puller | Sutter | P-97 |
| Borosilicate glass capillary | World Precision<br>Instrument | TW150-4 |
| 470 nm blue LED, mounted on a 10mm square<br>coolbase | LUXEON Rebel<br>LED | SR-05-B0040 |
| 700mA, buckpuck DC driver | LUXEON Rebel<br>LED | 3023-D-E-700 |
| Carclo 29.8° frosted 10mm circular beam optic-<br>integrated legs | LUXEON Rebel<br>LED | 10413 |
| 50mm square × 25mm high alpha heat sink | LUXEON Rebel<br>LED | N50-25B |
| Hand-held optical meter | Newport | 1918-R |
| BASIC Stamp2.0 microcontroller | Parallax | N/A |
| Digidata 1440 | Molecular Devices | https://www.moleculardevices.com |
| Multiclamp 700A Amplifier | Molecular Devices | https://www.moleculardevices.com |
| T-type thermal coupler | Omega | 5SRTC-TT-T-36-36 |
| Thermometer | Physitemp | BAT-10 |

### LEAD CONTACT AND MATERIALS AVAILABILITY

Requests for resources and reagents should be directed to and will be fulfilled by the Lead Contact, Yang Xiang.

### DATA AND CODE AVAILABILITY

All data and MATLAB code used for analysis and figure generation are available upon reasonable request.

### EXPERIMENTAL MODEL AND SUBJECT DETAILS

All fly stocks were raised in low yeast brown food at 25 °C and 60 % humidity in a 12 h: 12 h light dark cycle unless otherwise stated. *ppk-Gal4* (Han et al., 2012), *ppk-LexA* (Gou et al., 2014), *ppk-CD4-tdGFP* (Han et al., 2011)*, ppk-CD4-tdTomato* (Han et al., 2011), *ppk-EGFP* (Grueber et al., 2003), *19-12-Gal4* (Xiang et al., 2010), *UAS-CD4-tdGFP* (Han et al., 2011), *UAS-CD4-tdTomota* (Han et al., 2011), *UAS-mCD8-GFP* (Xiang et al., 2010), *UAS-Kir2.1* (Yang et al., 2014), *UAS-ChR2::EYFP* (Hwang et al., 2007), *UAS-TrpA1-B* (Gu et al., 2019), *UAS-AdoR* (Dolezelova et al., 2007) were described previously. *Repo-Gal4* (BL#7415), *NompC-Gal4* (BL#36361), *NompC-LexA* (BL#52241), *UAS-EYFP* (BL#6659), *UAS-EGFP* (BL#5430, BL#5431)*, UAS-GCaMP6s* (BL#42746, BL#42749)*, UAS-ChR2* (BL#52258), *UAS-CsChrimson::mCherry* (#BL82181), *UAS-Parvalbumin* (BL#25030), *UAS-Ras85D.N17* (BL#4845), *UAS-ras.N17* (BL#4846), *UAS-Ras85D.V12* (BL#4847, BL#64195, *UAS-TNT* (BL#28838), *AdoR* deletion mutant (*AdoR^-/-^*, BL#84447), *AdoR-t2A-Gal4* (BL#86138) were from Bloomington Drosophila Stock Center. *LexAop2-myr-GCaMP6s* was a gift from Dr. Marc Freeman (Oregon Health and Science University). All detailed cross information are provided in Supplementary Data.

### METHOD DETAILS

#### *In vivo* two-photon laser axotomy

Axotomy was performed at 80-86 hours after egg laying (AEL). We measured the axon regrowth between 24 hpa and 48 hpa time points as regeneration. We followed the previous protocol with several modifications (Song et al., 2012). Briefly, a larva was anesthetized with sevofluorane (Patterson Veterinary, NDC-14043-110-25) vapor for 3 min with dorsal side up. Axotomy was achieved using the bleaching function of a Zeiss LSM 7 MP two-photon microscope by localized axon injury (laser spot is ∼1.5 µm in diameter). We found 910 nm wavelength worked well for axotomy. Axons were severed at a point around half of the distance between the soma of C3da or C4da neurons and the soma of bipolar dendrite (BD) neurons (around 40-70 µm to the soma of C3da or C4da neurons). Following axotomy, larvae were recovered on damp Kimwipe paper and then were transferred to recovery vials containing regular brown food (for axon regeneration, Ca^2+^ imaging and electrophysiology) or white grape juice agar plates (for optogenetic induction of axon regeneration).

#### *In vivo* axon regeneration analysis

Quantitative analysis was performed as described previously (Song et al., 2019; Song et al., 2012). Briefly, the proximal axon length of C3da or C4da neurons were measured as **L_1_** at 24 hpa and **L_2_** at 48 hpa. Regeneration Length (**L_2_-L_1_**) is the increase of proximal axon length. The distances between the soma of C3da or C4da neurons and the soma of BD neurons were also measured as **BD_1_** at 24 hpa and **BD_2_** at 48 hpa to account for larval body size expansion during this period. Regeneration Index is calculated as: **L_2_/BD_2_**-**L_1_/BD_1_**.

#### Electrophysiology

Extracellular recording of spontaneous neuronal activity of C4da or C3da neurons was performed as described previously (Xiang et al., 2010). The *ex vivo* fillet preparation from 3^rd^ instar larvae was prepared at 24 hpa in external saline composed of (in mM): NaCl 120, KCl 3, MgCl_2_ 4, CaCl_2_ 1.5, NaHCO_3_ 10, trehalose 10, glucose 10, TES 5, sucrose 10, HEPES 10. Osmolality was 305 mOsm/kg. pH was adjusted to 7.25 by NaOH. The larval fillet preparation was mounted on a Zeiss Axio Examiner.D1 microscope with a water immersion objective lens (Objective W N-Achroplan 40x/0.75). Gentle negative pressure was applied to trap the neuronal cell body in a recording pipette (5 µm tip opening; 1.5–2.0 MΩ resistance) filled with the external saline. Recordings were performed with a Molecular Devices 700A amplifier, and the data were acquired with a Molecular Devices 1440A Digidata and Clampex 10.6 software (Molecular Devices). Extracellular recordings were obtained in voltage clamp mode with a holding potential of 0 mV, a 2 kHz low-pass filter and a sampling frequency of 20 kHz. Burst firing events were detected by the ‘Burst Analysis’ function in Clampfit 10.6 and were defined as a cluster of 5 or more consecutive action potentials, with an inter-spike interval of adjacent two action potentials less than 100 ms. The number of burst firing events divided by total recording duration (minute) was quantified as the frequency of burst firing events (bursts/min).

#### GCaMP imaging

Ca^2+^ signals of sensory neurons and ensheathing glia were measured by GCaMP6s imaging in larval fillet preparation as described previously (Gu et al., 2019) and in the same external saline used for electrophysiology. Time-lapse imaging was performed using the Zeiss LSM 700 confocal microscope, under a water immersion objective lens (Objective W Plan-Apochromat, 20x/1.0). Frame scan rate is 0.97 Hz. GCaMP signals were obtained from regions of interest (ROI) from soma, dendrite or axon for neurons or from glial branches. They are then corrected for horizontal drifting with ImageJ slice alignment plugin before analysis. Non-axotomized C3da or C4da neurons imaged from the same larvae were used as control for axotomized C3da or C4da neurons. Glia that ensheath the non-axotomized neurons from the same larvae were used as control for glia that ensheath the axotomized neurons. For imaging spontaneous Ca^2+^ signals of C4da neurons after dendrite severing, a primary dendrite branch of C4da neuron was severed by two-photon microscope laser, similar to axotomy, and GCaMP signals of C4da neurons were measured at 24 hours after dendrite injury. For imaging spontaneous Ca^2+^ signals of C4da neurons after UV-C treatment (Babcock et al., 2009), larvae were mounted dorsal side up with double-sided tape on a cover slide and mounted in an ultraviolet crosslinker (Spectroline, XL-1000). 20 mJ/cm^2^ 254 nm UV-C light was delivered. After treatment, larvae were recovered in regular food and GCaMP signals of C4da neurons were measured at 24 hours after UV-C illumination. For imaging ChR2-expressing C3da neuron responses to blue light stimulation, the average fluorescence signals of soma from 10 s before light stimulation was set as **F_0_**. The peak fluorescence signal during light stimulation was set as **F**. The response was quantified as **ΔF**/**F_0_**=(**F**-**F_0_**)/**F_0_**\*100%. For imaging TrpA1-B-expressing ensheathing glial responses to thermogenetic stimulation, 3^rd^ instar larval fillet was prepared in a customized silicone elastomer perfusion chamber. Pre-heated hot water (46 °C) was perfused through the inner cavity of the stainless-steel tube embedded in the perfusion chamber to apply heat stimulation. The saline temperature in the chamber was monitored by a T-type thermal coupler (Omega 5SRTC-TT-T-36-36) connected with a thermometer (Physitemp BAT-10), and recorded by the Clampex software (Molecular Devices). It took 40 to 60 s to elevate from room temperature (∼20 °C) to 30 °C. Temperature was then held at a value between 30 °C and 34°C for 2 min. Peak GCaMP signals of glial branch during this 2-min time-window were calculated as **F**. Average GCaMP signals in a 1-minute time-window before heating were calculated as **F_0_**. Glial responses were quantified as **ΔF**/**F_0_** =(**F**-**F_0_**)/**F_0_**\*100%. Larvae were raised at 21°C, a temperature point below the TrpA1-B activation threshold (Gu et al., 2019). For imaging C4da and C3da neuronal responses to thermogenetic activation of ensheathing glia, heating process was performed similarly to imaging TrpA1-B-expressing ensheathing glial responses. Temperature was held at a value between 30 °C and 34°C for 2 min. Peak somatic GCaMP signals during this 2-min time-window were calculated as **F**. Average GCaMP signals in a 1-min time-window before heating were calculated as **F_0_**. Neuronal responses were quantified as **ΔF**/**F_0_**=(**F**-**F_0_**)/**F_0_**\*100%. For imaging C4da neuron responses to optogenetic stimulation of ensheathing glia, average somatic GCaMP6s signals from 10 s before light stimulation were set as **F_0_**, and peak somatic GCaMP6s signals during light stimulation were set as **F**. The response was quantified as **ΔF**/**F_0_**=(**F**-**F_0_**)/**F_0_**\*100%. 555 nm green light pulses (500 ms on, 500 ms off, 5.0 mW/mm^2^) were repeated 10 times to stimulate glia. Larvae were raised in food containing 400 µM all-trans retinal (ATR, Sigma, R2500) at 25 °C in constant dark. For imaging responses of C4da and C3da neurons to adenosine, L-glutamate and D-serine, a 5-min time-window before and after drug perfusion was used to determine the frequency of neuronal Ca^2+^ spikes. For imaging ensheathing glial responses to AITC, average GCaMP signals of glial branches from 1min before AITC application was set as **F_0_**, and average GCaMP signals of glial branches from 2 min to 3 min after AITC application was set as **F**. The response was quantified as **ΔF**/**F_0_**=(**F**-**F_0_**)/**F_0_**\*100%.

#### Analysis of Ca^2+^ spike frequency

Time-lapse GCaMP signals (**F**) from C4da or C3da somatic regions were normalized to average intensity (**F_0_**) of each sample by (**F**-**F_0_**)/**F_0_**\*100%. After normalization, we used the Findpeaks function in Matlab to find local maxima to isolate Ca^2+^ spike peaks from background. GCaMP signals from non-axotomized C4da neurons also show weak Ca^2+^ spikes with small amplitude. To rule out these weak spikes and isolate strong spikes, we calculated standard deviation (**σ**) of GCaMP signals in each non-axotomized C4da neuron somatic regions, then calculated the mean value of **σ** (defined as **σ’**). We set minimum peak amplitude parameter in Findpeaks as three folds of **σ’** (3**σ’**) (Figure S4A), and only amplitude larger than 3**σ’** is considered as a Ca^2+^ spike. After acquiring the position, number and amplitude of each spike, we divided the number of spikes by the time duration to calculate Ca^2+^ spike frequency (Ca^2+^ spikes/min). Based on these criteria, the amplitudes of all Ca^2+^ spikes from non-axotomized C4da neurons were less than 25% (Figure S4D and S4E). In contrast, the amplitudes of around 60% of Ca^2+^ spikes from axotomized C4da neurons were larger than 25% (Figure S4C).

#### Optogenetic stimulation of axon regeneration *in vivo*

The setup for optogenetic stimulation of free moving larvae was modified from the previous report (Kaneko et al., 2017). Larvae were raised in food containing 400 µM ATR at 25 °C in constant dark in the fly incubator (Tritech research, DT2-MP-47L). At 80-82 hours AEL, early 3^rd^ instar larvae were subject to laser axotomy. Larvae were allowed to recover for 6h in darkness. Next, we transferred larvae to a 35 mm petri dish plate containing 1 mL solidified transparent grape agar. The plate was covered with transparent film. The formula of the agar plate was: 2.4 g agar, 2.6 g sucrose, 20 mL white grape juice, 2 ml 95% ethanol, then add ddH_2_O to 100 mL final volume. White grape juice was chosen to facilitate light penetration. To supplement ATR, grape agar was melted by microwave, and ATR was added to a final concentration of 400 µM after cooling down to 55 °C. The ATR-containing grape agar plates were freshly prepared, stored at 4 °C in dark, and used within 1-2 days. 470 nm blue LED (LUXEON Rebel LED, SR-05-B0040) was mounted on a 10 mm square coolbase and 50 mm square ×25 mm high alpha heat sink (LUXEON Rebel LED, N50-25B), set under circular beam optic with integrated legs (LUXEON Rebel LED, 10413) for even light illumination, and finally placed over the grape juice agar plate for optogenetic stimulation. The light pattern was programmed with BASIC Stamp 2.0 microcontroller and buckpuck DC driver (LUXEON, 700 mA, externally dimmable, 3023-D-E-700). Light intensity (1.4 mW/mm^2^ 470 nm blue light) was measured by a power meter (Newport, 1918-R). Free moving larvae in the grape juice agar plate were continuously stimulated with three different parameters that elicited burst, semi-burst, and tonic firing in C3da neurons (Figure 4D-4E). At 24 and 48 hpa, axon lengths were imaged and measured by confocal microscope and regeneration index was calculated.

#### Immunostaining of *Drosophila* larvae

For immunostaining with the antibody against dpERK, 3^rd^ instar larvae were dissected in ice-cold external saline used for electrophysiology. Between each of the following treatment, fillets were washed 4 times of 15 min, each with washing buffer containing 0.3% Triton X-100 (Fluka, BP151) in phosphate buffered saline (PBS, Sigma). Larvae were firstly fixed with PBS containing 4% paraformaldehyde (PFA, Sigma) for 30 min. Then larvae were treated with blocking buffer for 1h at room temperature. The blocking buffer was PBS containing 0.3% Triton X-100, 5% donkey serum (Sigma, D9663), and 0.1% bovine serum albumin (Sigma). Next, larvae were incubated in rabbit anti-Phospho-p44/42 Erk1/2 (Thr202/Tyr204) primary antibody (Cell signaling, rabbit, #4370S, RRID:AB2315112, 1:100 in blocking buffer) for 18 hours at 4°C, and then in the goat anti-rabbit Alexa 647-conjugated secondary antibody (Jackson immunoresearch, 111-605-144, RRID:AB_2338078, 1:500 in blocking buffer) for 2 hours at room temperature. After final wash, larvae were mounted using the anti-fade mountant (Thermo Fisher, P36961) prior to confocal imaging. For dpERK immunostaining after AITC stimulation, larvae were treated with external saline containing 100 µM AITC for 15 min before proceeding with the above immunostaining protocol.

#### Intravitreal injection and optic nerve crush

Intravitreal viral or drug injection, optic nerve crush and RGC axon labeling were performed as previously described (Park et al., 2008). Briefly, under anesthesia, rAAV2 (2 µL), adenosine (10 µM in 2 µL PBS, Sigma), PBS (2 µL) or Alexa 555-conjugated cholera toxin beta subunit (CTB-555, 1 mg/ml; 2 µL; Thermo Fisher) was injected intravitreally with a fine glass micropipette connected to the Hamilton syringe using plastic tubing. The position and direction of the injection were well controlled to avoid lens damage. For optic nerve crush in anesthetized animals, the optic nerve was exposed intraorbitally and crushed using a pair of Dumont #5 forceps (Fine Science Tools) for 5 s at approximately 1 mm behind the optic disc at 14 days after AAV2 injection. Adenosine or PBS was injected intravitreally for 5 times every 3 days after optic nerve crush. CTB-555 injection was performed 2 days before euthanasia to trace regenerating RGC axons. At the final step of the experiment procedure, both retina and the optic nerve were dissected out and post-fixed in 4% PFA overnight at 4°C for further axon regeneration and neuronal survival quantifications. rAAV2-hSyn-Adora2b and rAAV2-hSyn-PTEN-shRNA were packaged by the virus core in CEBSIT (Center for Excellence in Brain Science and Intelligence Technology, Chinese Academy of Sciences, Shanghai, China). All virus used had titers > 1 × 10^13^.

#### Optic nerve dehydration and clearing

Dehydration and clearing of optic nerves were performed following previous reports (Belle et al., 2017; Erturk et al., 2012). Briefly, fixed optic nerves were first dehydrated in incremental concentrations of tetrahydrofuran (TFH, 50%, overnight; 80%, 1 hour and 100%, 2 hours, v/v % in distilled water, Sigma) in amber glass bottles with a silicon-coated cap (Thermo Fisher). Incubations were performed on a shaker at room temperature. Then the nerves were incubated in Dichloromethane (DCM, Sigma) until the nerves sinks at the bottom (5 min to 1 hour max), then in DiBenzyl Ether (Sigma) until the sample is clear (20 min to 2 hours). The nerves were protected from light during the whole process to avoid potential photo bleaching of the fluorescence.

#### Analysis of RGC axon regeneration

The cleared whole-mount optic nerves were imaged with a 10x objective lens on a Nikon C2 confocal microscope equipped with a motorized stage and Nikon software. For each optic nerve, we used the Z-stack function to acquire a stack of 8.17-µm-thick slices, with 60% overlap between adjacent slices. The slices were then stitched to obtain an image of the whole-mount nerve. Quantification of regenerating axons in the optic nerve was performed as previous described (Park et al., 2008; Wang et al., 2018). Specifically, every 10 consecutive slices were Z-projected with maximum intensity to generate a series of Z-projection images of 81.7-µm-thick optical sections. At every 250-µm interval from the crush site, we counted the number of individual CTB-labeled axons and measured the width of the nerve at each optical section site. Both numbers were used to calculate the number of axons per micrometer of nerve width, which was then averaged over all optical sections as reported previously (Wang et al., 2018). Σ**ad**, the total number of **a**xons extending distance **d** in a nerve with a radius of **r**, was estimated by summing over all optical sections with a thickness of **t** (81.7 µm): Σ**ad** = π**r**^2^ × (average axons/µm)/**t**.

#### Analysis of RGC survival

Following overnight incubation in 30% sucrose, the whole retina was first incubated in optimal cutting temperature (OCT) compound for cryosection and then was sectioned for staining. The cryosections were washed with PBS for 3 times of 15 min, and then blocked in PBS containing 2% bovine serum albumin and 0.1% Triton X-100 for 1 hour. We immunostained the retina neurons with mouse anti-Tuj1 (tubulin beta 3) primary antibody (BioLegend, #801202, RRID: AB_10063408) and donkey anti-mouse Alexa 488-conjugated secondary antibody (abcam, #150109) to examine the number of surviving RGCs after injury. Fluorescent images were acquired with a 20x objective on a Zeiss Imager M2 microscope. RGC numbers were counted from at least three 20 µm thick retinal sections per retina (approximately 500-800 µm per section) using the cell counter plugin from ImageJ software. We calculated the linear density of Tuj1 positive cells in the ganglion cell layer (GCL) and normalized these counts to a standard intact control non-injured retina, as reported previously (Park et al., 2008; Wang et al., 2018).

#### Quantification and statistics

Data were represented as mean ± S.E.M. in point graphs. All statistical analyses and graph making were performed using GraphPad Prism 9 (GraphPad Software). We did not pre-determine sample sizes and our sample sizes are similar to those reported in previous publications (Song et al., 2019; Song et al., 2012). Mann-Whitney rank-sum test or Student’s *t*-test was performed for comparisons between two groups of samples. Kruskal-Wallis statistic test followed by Dunn’s multiple comparison tests or one-way ANOVA followed by Bonferroni’s, Dunnetts’s or Tukey’s multiple comparison tests were performed for comparisons among three or more groups of samples. Two-tailed unpaired *t*-tests were performed with Welch’s correction to adjust for unequal variance. Two-sided Wilcoxon matched-pairs signed-rank test was used to analyze data that did not follow normal distribution. The sample sizes of ‘n’ are provided in the figure legends. All detailed statistical analyses are provided in Table S1.

## SUPPLEMENTAL VIDEO LEGEND

**Video S1. An example video for GCaMP6s imaging showing that glial thermogenetic stimulation activates non-axotomized C4da neurons.** A single C4da neuron is labeled by *ppk-LexA>LexAop2-myr-GCaMP6s*. TrpA1-B was expressed in glia. Scale bar, 20 µm.

**Video S2. An example video for GCaMP6s imaging showing that glial thermogenetic stimulation did not activate non-axotomized C3da neurons.** A single C3da neuron is labeled by *NompC-LexA>LexAop2-myr-GCaMP6s*. TrpA1-B was expressed in glia. Scale bar, 20 µm.

**Video S3. An example video for GCaMP6s imaging of non-axotomized C4da neurons.** A single C4da neuron is labeled by *ppk-LexA>LexAop2-myr-GCaMP6s*. Scale bar, 20 µm.

**Video S4. An example video for GCaMP6s imaging of axotomized C4da neurons at 24 hpa.** A single axotomized C4da neuron is labeled by *ppk-LexA>LexAop2-myr-GCaMP6s*. Each quantified Ca^2+^ spike peak is highlighted by beep sound. Scale bar, 20 µm.

**Video S5. An example video for simultaneous extracellular electrophysiological recording and GCaMP6s imaging of axotomized C4da neuron at 24 hpa.** A single axotomized C4da neuron is labeled by *ppk-LexA>LexAop2-myr-GCaMP6s*. The soma region is defined as ROI for GCaMP6s signals. Scale bar, 20 µm.

## SUPPLEMENTAL TABLE LEGEND

**Table S1. Detailed fly crossing information, sample size number and statistics in each main figure panel and supplemental figure panel.**

## Notes

### Competing Interest Statement

The authors have declared no competing interest.

